# The drug-induced phenotypic landscape of colorectal cancer organoids

**DOI:** 10.1101/660993

**Authors:** Johannes Betge, Niklas Rindtorff, Jan Sauer, Benedikt Rauscher, Clara Dingert, Haristi Gaitantzi, Frank Herweck, Kauthar Srour-Mhanna, Thilo Miersch, Erica Valentini, Veronika Hauber, Tobias Gutting, Larissa Frank, Sebastian Belle, Timo Gaiser, Inga Buchholz, Ralf Jesenofsky, Nicolai Härtel, Tianzuo Zhan, Bernd Fischer, Katja Breitkopf-Heinlein, Elke Burgermeister, Matthias P. Ebert, Michael Boutros

## Abstract

Patient derived organoids resemble the biology of tissues and tumors, enabling *ex vivo* modeling of human diseases from primary patient samples. Organoids can be used as models for drug discovery and are being explored to guide clinical decision making. Patient derived organoids can have heterogeneous morphologies with unclear biological causes and relationship to treatment response. Here, we used high-throughput, image-based profiling to quantify phenotypes of over 5 million individual colorectal cancer organoids after treatment with more than 500 small molecules. Integration of data using a joint multi-omics modelling framework identified organoid size and cystic vs. solid organoid architecture as axes of morphological variation across organoids. Mechanistically, we found that organoid size was linked to IGF1 receptor signaling, while a cystic organoid architecture was associated with an LGR5+ stemness program. Treatment-induced organoid morphology reflected organoid viability, drug mechanism of action, and was biologically interpretable using joint modelling. Inhibition of MEK led to cystic reorganization of organoids and increased expression of LGR5, while inhibition of mTOR induced IGF1 receptor signaling. In conclusion, we identified shared axes of variation for colorectal cancer organoid morphology, their underlying biological mechanisms, and pharmacological interventions with the ability to move organoids along them. Image-based profiling of patient derived organoids coupled with multi-omics integration facilitates drug discovery by linking drug responses with underlying biological mechanisms.

## Introduction

Colorectal cancer is globally the third most common cancer and the second leading cause of cancer related death.^1^ Patients with advanced and metastatic disease are usually treated with chemotherapy and antibody therapies, however, with current treatment, tumors may continue to progress and prognosis remains poor.^2^ Tumor plasticity, as well as the stemness of neoplastic cells have been proposed as major factors in treatment resistance and tumor progression under antineoplastic therapies.^3,4^ However, mechanisms behind these tumor cell states and drugs modulating or targeting them are not well understood.

Patient derived organoids (PDOs) are stem cell derived 3D tumor models that can be efficiently established from (colorectal-) cancer and normal tissues.^5–7^ Organoid isolation from human primary tumors and metastases^5,8^ has enabled the establishment of living biobanks.^6,7,9^ Notably, patient derived organoids have been shown to represent their origin’s molecular features and morphology,^6–8,10^ enabling functional experiments such as drug testing *ex vivo*.^7,9,11–16^ As a consequence, organoids are an attractive model system, as they combine the modeling capacity of patient derived xenografts with the scalability of adherent in vitro cell lines.

Image-based profiling is a high-throughput microscopy-based methodology to systematically measure phenotypes of *in vitro* models. When combined with chemical or genetic perturbations, image-based profiling is a powerful approach to gain systematic insights into biological processes, for instance in drug discovery and functional genomics research^17–19^. Image-based assays have been used to screen large libraries of small molecules to identify potential drug candidates, to analyze a drug’s mode of action, or to classify drug-gene interactions by cell-morphology ^20–23^. Performing large image-based profiling experiments of organoids has been, however, a biological, technical and computational challenge^24–26^. Consequently, the morphological heterogeneity of patient-derived cancer organoids between and within patient donors, their diverging behaviors upon pharmacological perturbation, as well as the underlying mechanisms of cancer organoid morphology are not yet systematically understood.

Here we report a large-scale image-based phenotyping study of patient derived cancer organoids to understand underlying factors governing organoid morphology. Colorectal cancer organoids from 11 patients were treated with more than 500 experimental and clinically used small molecules at different concentrations. We systematically mapped the morphological heterogeneity of patient derived organoids and their response to compound perturbations from more than 3,700,000 confocal microscopy images. We found that the resulting landscape of organoid phenotypes was mainly driven by differences in organoid size, viability and cystic vs. solid organoid architecture. Using multi-omics factor analysis for integrating organoid morphology, size, gene expression, somatic mutations and drug activity, we identified biological programs underlying these phenotypes and small molecules that modulate them.

## Results

### Image-based profiling captures the morphological diversity of patient-derived cancer organoids

To better understand the diversity of organoid phenotypes, drug-induced phenotypic changes and the underlying factors driving them, we generated PDOs from 13 colorectal cancer patients representing different clinical stages and genotypes (Supplemental Fig. S1a-d, Supplemental Tables 1 and 2). We performed image-based profiling at single organoid resolution with 11 organoid lines (Fig. 1a) using small molecules targeting developmental pathways, protein kinases (464 compounds at a single 7.5µM concentration), as well as small molecules in clinical use (63 compounds in 5 concentrations, Supplemental Fig. S2a-c). After three days of culture and four days of pharmacological perturbation in 384-well plates, organoids were subsequently stained with fluorescent markers for actin (Phalloidin), DNA (DAPI), and cell permeability (DeadGreen) to capture their morphology with high-throughput confocal microscopy. We projected the 3D image data onto a 2D plane, segmented organoids and calculated morphological profiles for each organoid spanning 528 phenotypic features (such as dye intensity, texture, and shape) that were subsequently reduced into 25 principal components representing 81% of morphological variance (Supplemental Fig. S2c).

**Fig. 1:**
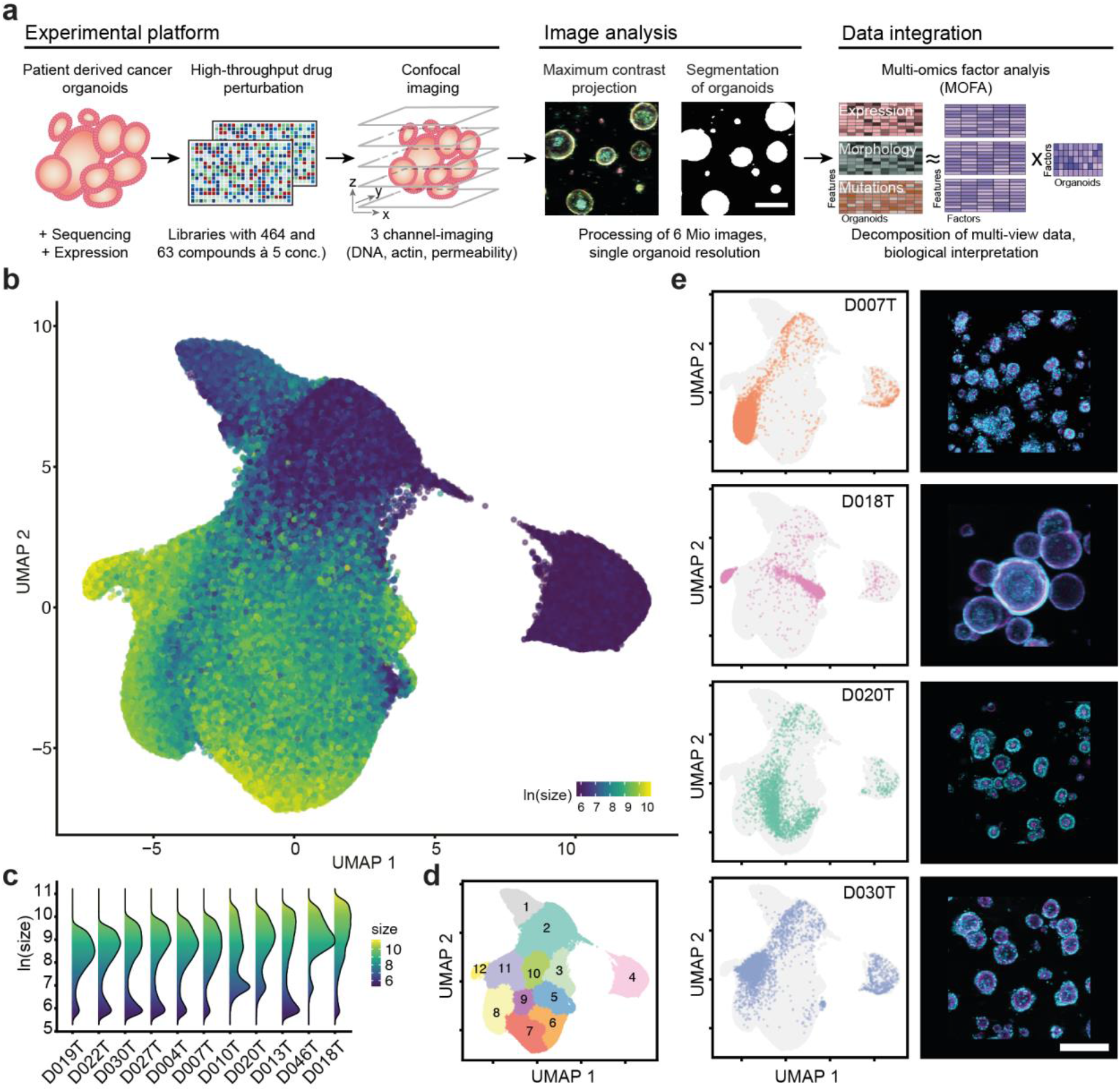
Image-based profiling captures the phenotype diversity of patient derived cancer organoids. **a**, Schematic overview of experiments: Organoids were isolated from endoscopic biopsies from patients with colorectal cancer. Organoids were dissociated and evenly seeded in 384-well plates before perturbation with an experimental (464 compounds) and a clinical compound library (63 compounds at 5 concentrations each, 842 perturbations across both libraries). After treatment, high-throughput fluorescence microscopy was used to capture the morphology of organoids. The multi-channel (DNA, beta-actin, cell permeability) 3D imaging data was projected, segmented, and phenotype features were extracted to quantify potential drug-induced phenotypes. Untreated organoid morphology, organoid size and drug activity scores were integrated with mRNA expression and mutation data in a Multi-Omics Factor Analysis (MOFA). **b**, Uniform Manifold Approximation and Projection (UMAP) of organoid-level features for a random 5% sample out of approximately 5.5 million organoids. The same sample is used for visualizations throughout the figure. Color corresponds to the log-scaled organoid area (dark blue: minimum size, yellow: maximum size). **c**, organoid size distribution across organoid lines **d**, UMAP representation of DMSO treated and drug treated organoids. Graph-based clustering of organoids by morphology with 12 resulting clusters. **e**, UMAP embeddings of selected organoid lines (baseline state = 0.1% DMSO control-treated organoids) representing different morphological subsets, grey background consists of randomly sampled points. Depicted are representative example images for each organoid line (right, cyan = DNA, magenta = actin, scale-bar: 200µm).

To visualize the heterogeneity of colorectal cancer organoids and treatment induced changes across and within PDO lines, we embedded features of approximately 5.5 million profiled organoids using uniform manifold approximation and projection (UMAP, Fig. 1b, Supplemental Fig. 2d-f). Organoids showed a characteristic two-component log-normal mixture distribution of organoid size within most lines, with one component containing small organoids and another component containing larger organoids with varying, organoid line specific, reproducible average size (Fig. 1c, Supplemental Fig. S2g-h). This size distribution likely resulted from intrinsic differences in cellular size and growth rate accumulating throughout the course of the experiment in multicellular organoids. Next, we performed graph-based clustering on this embedding to describe the landscape, resulting in 12 clusters (Fig. 1d). Organoid lines within the embedding were located in characteristic clusters, with organoid size and organoid architecture as primary organizing factors (Fig. 1e). For example, organoid line D018T had the largest median organoid size within the dataset and a cystic organoid architecture with a single central hollow lumen and monolayer of surrounding cells. In contrast, D020T organoids had a solid architecture and smaller median size. In most cases, organoid lines had two areas of main density, with one of them in clusters 2, 3 or 4, reflecting the previously mentioned bimodal size distribution. When comparing drug-treated organoids to baseline organoids treated with the solvent control (DMSO), no clear separation of groups was apparent, suggesting that organoid morphology was distributed on a continuum of phenotypes spanning perturbed and unperturbed conditions of our experiment (Supplemental Fig. S2i). In summary, image-based profiling of patient derived colorectal cancer organoids showed strong morphological heterogeneity with donor dependent differences in size and organoid architecture.

**Fig. 2:**
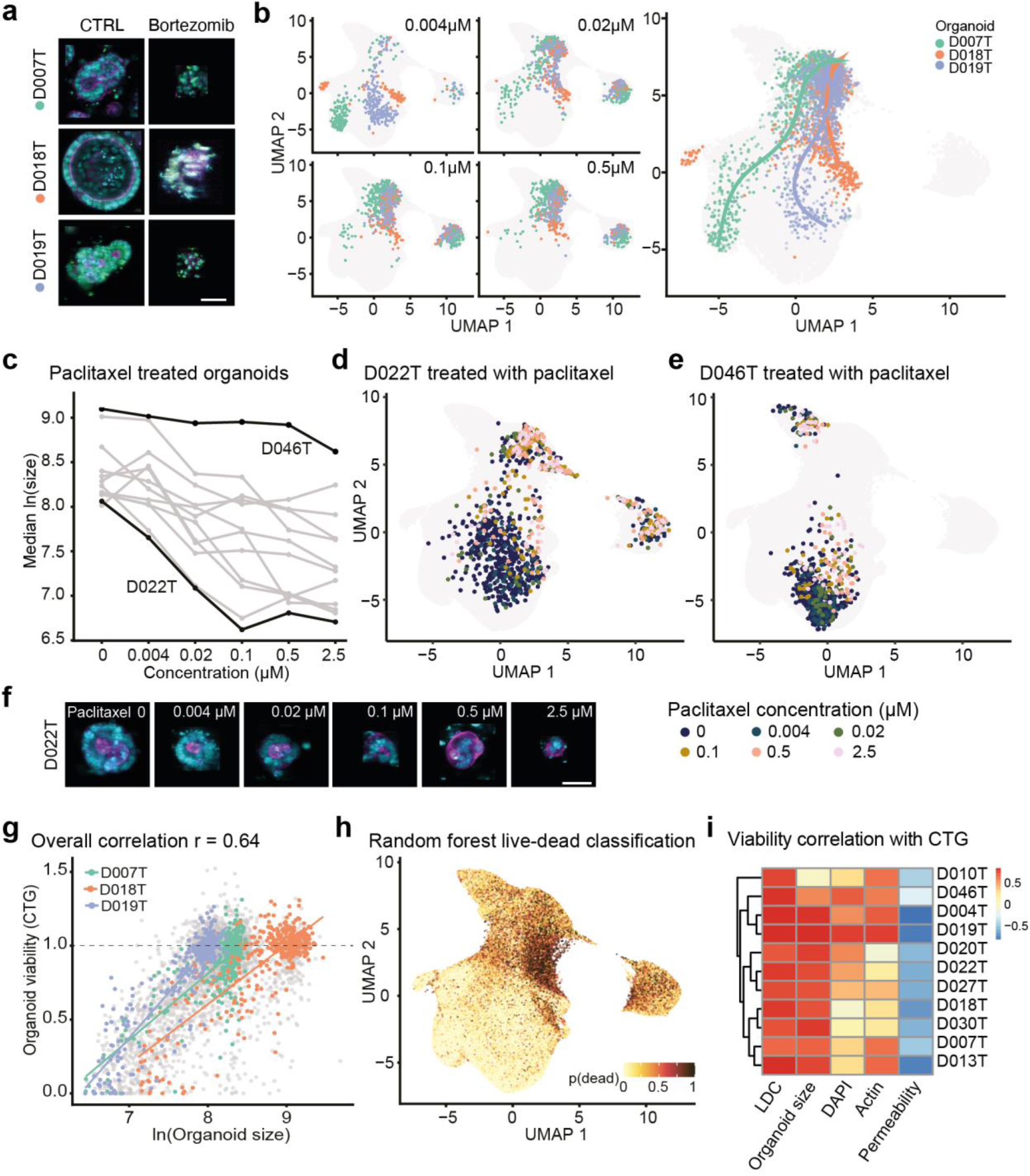
Organoid phenotype-profiles capture organoid viability. **a**, Representative example images of negative (0.1% DMSO) and positive control treated organoids (2.5µM bortezomib, cyan = DNA, magenta = actin, yellow = cell permeability; representative images were selected and embedded in black background; scale bar: 50µm). **b**, Dose-dependent-trajectory of bortezomib drug effect. UMAP of organoid morphology at different bortezomib doses and (right panel) dose-dependent trajectory for three representative organoid lines. During the principal curve fitting, trajectory inference excluded cluster 4, a set of measurements representing mostly dead organoid particles comprising ca. 5% of all imaging data. **c**, Dose-response relationship of organoid size and paclitaxel dose. D022T and D046T are highlighted as examples for responder/non-responder lines. **d**, UMAP of organoid morphology highlighting D022T organoids treated at different concentrations of paclitaxel. **e**, D046T organoids treated at different concentrations of paclitaxel. **f**, Example images of D022T organoids treated with paclitaxel. **g**, Association of organoid size of selected example organoid lines with viability determined by luminescence-based, ATP-dependent viability profiling with CellTiter-Glo (CTG), which was performed in parallel with imaging on a subset of drug treatments for benchmarking. **h**, UMAP visualization of viability predictions for organoids within our dataset, based on supervised machine learning of organoid viability using classifiers trained on positive (high-dose bortezomib and SN-38) and negative (DMSO) controls (live-dead classifiers, LDC). **i**, Association of LDC and example organoid features (size, DAPI, actin and permeability dye intensities) with benchmark CTG viability read out.

### Organoid phenotype profiles capture organoid viability

Drug-induced changes in cell viability are a fundamental readout in cancer drug discovery. Prompted by the observation that organoid size was a major factor determining the phenotype embedding, we hypothesized that small organoid size, which was seen across all donors, was at least partially the result of cell death within organoids and, more broadly, that phenotype data could be used to estimate organoid viability. To test this hypothesis, we chose bortezomib, a small molecule proteasome inhibitor with high *in vitro* toxicity, as well as SN-38 (active metabolite of irinotecan). Both small molecules led to dose dependent organoid death in all organoid lines (Fig. 2a). Analogous to pseudotime in single-cell gene expression analysis,^27^ we fitted dose-dependent trajectories of bortezomib (Fig. 2b) and SN-38 (Supplemental Fig. 3a). Starting from diverse baseline morphologies, increasing doses of these compounds led to a stepwise convergence on a final death-related phenotype, which corresponded to the areas with enrichment of small objects (clusters 2, 3 and 4 shown in Fig. 1d). Similarly, paclitaxel, a microtubule disassembly inhibitor, shifted the bimodal size distribution of organoids in a dose-dependent fashion (Supplemental Fig. S3b), while organoid count remained largely unchanged (Supplemental Fig. S3c). This effect, however, was organoid line-specific, as we observed a dose-dependent decrease in median organoid size in paclitaxel “responder” lines (e.g. D022T), while the size of other organoids remained unaffected (e.g. D046T, Fig. 2c-f). These observations suggested a link between organoid morphology, especially organoid size, with a loss of cell viability. To test the ability of organoid morphology to predict cell viability, we performed a luminescence-based, ATP dependent, cell viability assay (CTG) in parallel with imaging as a benchmark for drugs within the clinical cancer panel. We saw a strong association of CTG viability with organoid size (Fig. 2g), prompting us to test whether a more accurate prediction of organoid viability was possible by using all available imaging information (including organoid size). To this end, we trained random forest classifiers (live/dead classifiers, LDC) on individual organoid phenotype profiles to distinguish between negative and positive control treatments (DMSO, bortezomib and SN-38, Supplemental Fig. S3d-e). We observed robust classification performance when applied to sets of the same or unseen organoid lines (Supplemental Fig. S3f). As expected, when applying the classifier to the whole imaging dataset and visualizing predictions via UMAP, small organoids within previously identified clusters 2, 3 and 4 had the highest probabilities for death (Fig. 2h).

**Fig. 3:**
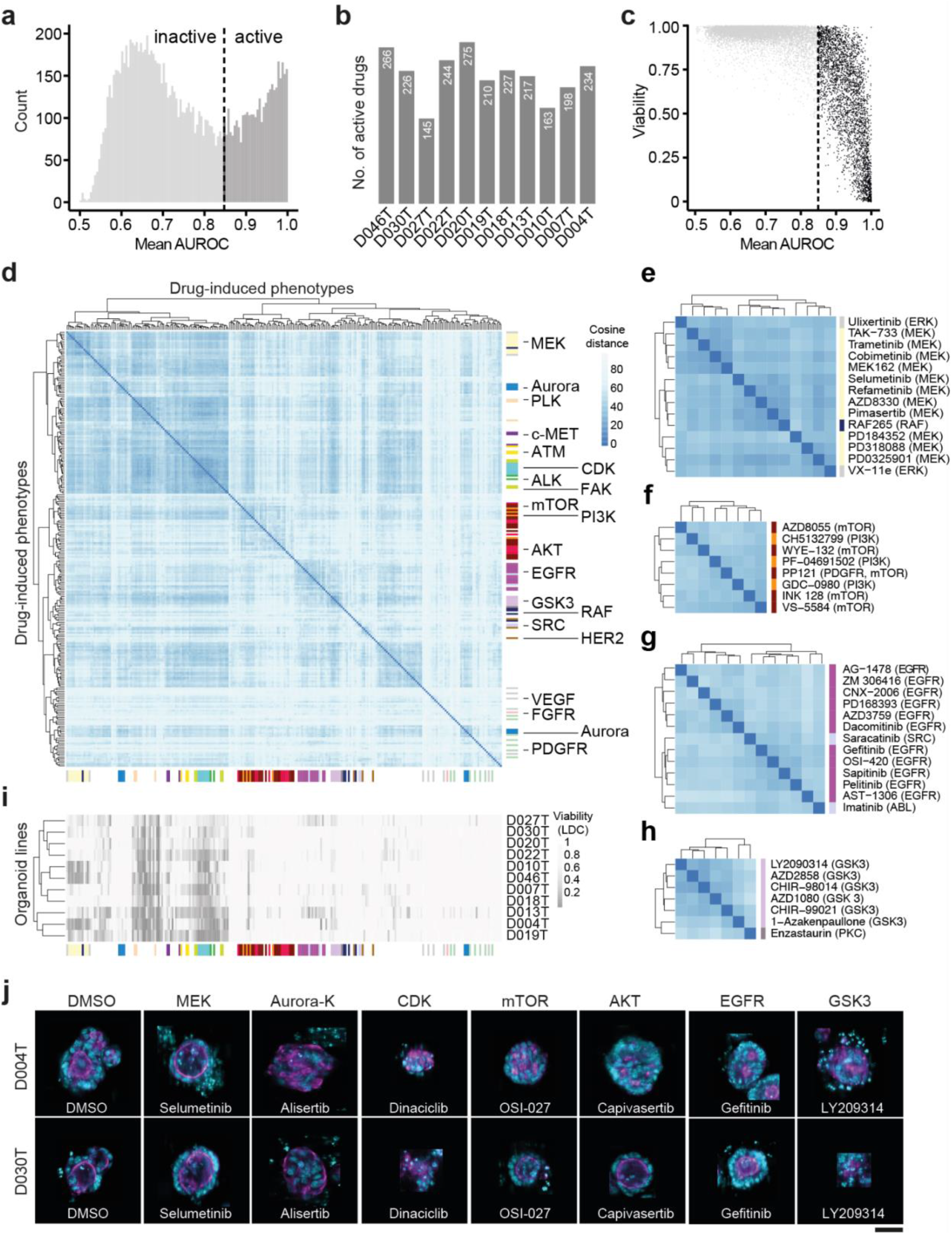
Drug-induced organoid phenotypes correspond to drug mechanism of action. **a**, Histogram of average model performance for each tested drug. For every tested drug and organoid line, logistic regression models were trained to distinguish negative control-treated organoids from drug treated organoids. A drug was considered “active” when it induced a phenotype that could be separated from DMSO control-phenotypes with a mean classification performance of >0.85 area under the receiver operating characteristic curve (AUROC). **b**, Number of active drugs per organoid line. **c**, Relationship between drug-induced viability change (predicted by LDC, compare Fig. 2) and general compound activity. **d**, Unsupervised clustering of drug effect profiles for active drugs. Distance between drug effect profiles was calculated using cosine similarity. Drug effect profiles were determined by fitting logistic regression models between treated and untreated organoids for each drug and line. PCA transformed morphology information was used as input features. Fisher’s exact test was used to identify enrichments of drugs annotated with the same drug target within the hierarchical clustering. Tested clusters had a minimum cluster size of 3 and were evaluated iteratively from the tree bottom to top. Colors on the side of the heatmap represent drug mechanisms of action. **e-h**, Zoom-ins of (d) showing clusters enriched for MEK (e), PI3K/mTOR (f), EGFR (g) and GSK-3 (h). **i**, Viability of drug-induced phenotypes in individual organoid lines as determined by supervised machine learning. The drugs were arranged on the x-axis in the same order as in (d). **j**, Organoids representative of selected drug-induced phenotypes. Images from organoid lines D004T and D030T were selected for each organoid phenotype, automatically cropped and embedded in black background. Cyan = DNA, magenta = actin; scale bar: 50µm.

The LDC predictions had the highest correlation with CTG based viability data (Fig. 2i), however, the association with organoid size was almost as strong in the majority of organoid lines (Fig. 2i, Supplemental Fig. S3g), while other simple features, such as DNA (DAPI), actin (phalloidin), and especially permeability (DeadGreen) intensity in isolation were less suitable to predict viability of organoids (Fig. 2i). We also noticed in ablation experiments that LDCs with incomplete access to channel information (i.e. only DAPI and phalloidin staining derived features were available for training and inference) showed, in some instances, classification accuracies almost as high as classifiers with access to complete data (Supplemental Fig. S3h). Finally, we observed examples of diverging results between LDC predictions and CTG read-outs. These included (1) the antifolate drug methotrexate and (2) a doxorubicin-induced artifact due to the strong red color of the compound. Methotrexate showed strong toxicity in almost all organoid lines in CTG based experiments but had no visible effect on organoid viability based on the LDC (Supplemental Fig. S2i-l). This discordance may be explained by non-lethal metabolic effects of methotrexate. In conclusion, basic features, such as median organoid size, as well as classification of texture and shape information from basic DNA and actin staining could predict organoid viability.

### Drug-induced organoid phenotypes correspond to drug mechanism of action

An advantage of image-based profiling over cell viability measurements in drug discovery is the ability to use the high dimensional drug-induced phenotype profiles to identify active but not necessarily lethal small molecules and estimate their mechanism of action by similarity-based clustering. To test whether this approach could be used in patient derived cancer organoids, we used a supervised learning approach for drugs within the KiStem library to identify drug effect profiles and group them by similarity. First, we trained logistic regression models to distinguish individual compound-treated organoids from unperturbed controls and defined the resulting normal vector between control- and treated organoid phenotypes as the drug effect profile. Next, we scored every logistic regression model’s ability to separate treated and untreated organoids to identify active treatments that induced a robust change in organoid morphology (area under the receiver operating characteristic, AUROC, ranging from 0.5 to 1). We considered treatments active when their classifiers’ performance exceeded an AUROC of 0.85 (Fig. 3a, 3b). Based on our observations, drug activity was necessary but not sufficient for a viability effect (Fig. 3c) as a fraction of drugs led to identifiable changes in organoid morphology (they were considered active drugs) but were not classified as lethal by our live/dead classifier (LDC) models.

To test whether active drugs systematically induced organoid phenotypes that were informative of mechanism of action, we assessed similarity by two different methods, (1) the cosine distance between concatenated drug effect profiles and (2) the Euclidean distance of averaged treatment-induced phenotypes (Fig. 3d-h, Supplemental Fig. S4a-c, S5a-c). While both methods were similar in terms of their ability to cluster drugs by mechanism of action, we proceeded with cosine distance clustering, as drug effect profiles did not only capture the direction of phenotype change, but were also linked to AUROC as a metric of drug activity that was scaled between 0.5 and 1. We observed a clustering of drugs by their specific mode-of-action, including inhibitors of MEK, aurora kinase, CDK, mTOR, AKT, EGFR or GSK3 (Fig. 3d). Small molecules with targets within the same signaling pathway also induced related morphologies, for example MEK inhibitors clustered with specific RAF- and ERK inhibitors (Fig. 3e) and AKT/ PI3K inhibitors were part of a cluster mainly containing mTOR targeting small molecules (part of the cluster is shown in Fig. 3f, whole cluster in supplemental Fig. S5a). The clustering also suggested additional mode-of-actions or off-target effects for well-described small molecules (Fig. 3g-h). For example, the PKC inhibitor enzastaurin was clustered with GSK3 inhibitors, substantiating a previously described interaction of enzastaurin with the alpha and beta subunits of GSK3^28,29^ (Fig. 3h). Of note, several drug-induced phenotypes were observable across most, or all tested organoid lines, but the majority of compound classes led to significant enrichments in drug profile vector clustering only in subsets or individual organoid lines (Supplemental Fig. S5d).

To assess whether morphological profiles of active drug treatments were primarily driven by differences in organoid viability, we compared LDC predictions with the phenotypic clustering (Fig. 3i). We observed a larger cluster of lethal treatments (including molecules targeting ATM, JAK, PLK, CDK). However, most clusters were caused by non-lethal phenotypes, including those induced by inhibitors of AKT, mTOR, EGFR or GSK3. Visual inspection of several phenotypes (Fig. 3j) revealed recurring drug target dependent morphologies. Most notably, MEK inhibitors led to reorganization towards a more cystic organoid architecture. Altogether, drug-induced phenotypes were capturing drug mode of action and were visible across most tested organoid lines.

### Multi-omics factor analysis identifies shared factors linking morphology, genomic data and drug activity

A limitation of image-based profiling experiments is that both unperturbed and drug-induced morphologies are challenging to interpret in terms of their underlying biology. Theoretically, in the presence of multiple *in vitro* models with both phenotype and genomic measurements, links between the two data modalities can be learned. Based on the observation that organoid morphology was distributed in a continuous space, we hypothesized that variation in organoid baseline morphology could be associated with differences in gene expression, mutations, as well as drug activity for the 11 cancer organoid lines in our sample (2 biological replicates each, 22 observations in total). To factorize the joint distribution of unperturbed organoid morphology, unperturbed organoid size, gene expression, selected somatic mutations, and drug activity, we performed multi-omics factor analysis (MOFA).^30^ MOFA is a matrix factorization method that decomposes a set of different measurements into a shared table of factors scoring each observed sample and a set of corresponding loading tables linking each factor to features in the set of original measurements.^30^. When trained with k = 3 factors, MOFA recovered factors explaining approximately 24-41% of variance across the different data modalities (Fig. 4a-b, Supplemental Fig. S6a-c). While gene expression, mutations and drug activity profiles for organoid lines contributed to all factors, factor 1 captured most variation in median organoid size (ca. 39%). In contrast, factor 2 was primarily capturing variation within untreated organoid morphology (ca. 16%) (Fig. 4a). Organoid lines D046T and D004T stood out as lines with the strongest score for factor 1, while organoid lines D018T and D013T had the strongest score in factor 2 (Fig. 4c). Organoids with high factor scores were in characteristic regions of the previously defined UMAP embedding (Supplemental Fig. S6d). Visual inspection of organoids revealed that organoid lines with a higher factor 1 score tended to be larger in size and organoids with high factor 2 score tended to have a more cystic organoid architecture based on manual classification. Analysis of gene expression data alone recovered patterns analogous to factor 1 and factor 2 (Fig. S6e-f). We could not identify interpretable morphological differences between factor 3 low and high organoids and focused our subsequent analysis on the first two interpretable factors generated by MOFA. In summary, MOFA identified factors within the dataset that explained variation between organoid lines across different data modalities, including organoid morphology and median size.

**Fig. 4:**
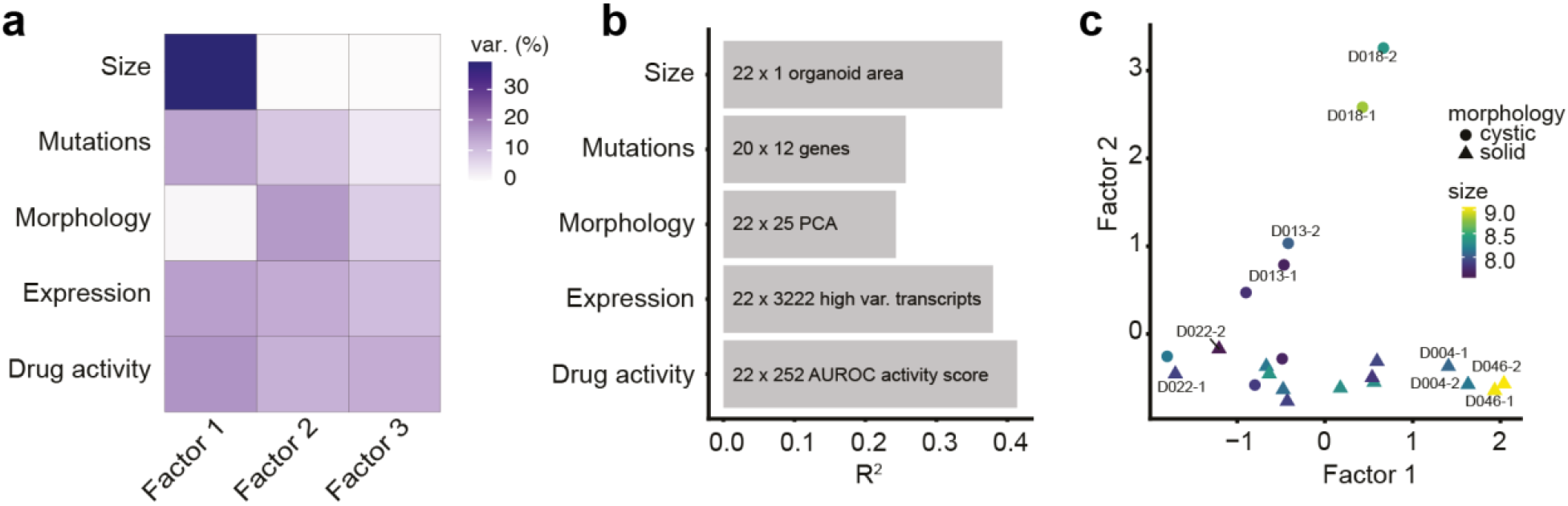
Multi-omics factor analysis identifies shared factors linking morphology, gene expression and drug activity. **a-b**, Variance decomposition of the MOFA model. Untreated organoid morphology, organoid size and drug activity scores were integrated with DNA sequencing and mRNA expression data. **a**, Percentage of variance explained by each factor in each data modality. **b**, Cumulative proportion of total variance explained by each experimental data modality within the MOFA model. **c**, Visualization of samples in factor space showing factors 1 and 2. Shown are two independent replicates for each organoid line. Organoid morphology (cystic vs. solid) as determined by visual inspection of DMSO phenotypes and organoid size (log-scaled organoid area) are represented by symbol shape and color, respectively.

### An LGR5+ stemness program is associated with cystic organoid architecture and can be induced by inhibition of MEK

A particularly strong recurring organoid phenotype was the presence of a cystic organoid architecture, seen in untreated D018T or D013T organoids and organoids treated with MEK inhibitors (Fig. 1e, 3f, 5a). MOFA showed that factor 2 represented this cystic organoid state. In the cystic state, organoids consisted of a monolayer of uniform cells lining a central spherical lumen with a pronounced actin cytoskeleton (Fig. 5b). We considered this phenotype related to organoid morphologies previously described in genetically engineered APC-/- or Wnt ligand treated intestinal organoids.^31–33^ To test if factor 2 in fact captured Wnt signaling and intestinal stem cell identity related gene expression programs, we performed gene set enrichment analyses (GSEA) for cell identity signatures previously identified in intestinal crypts and colorectal cancer.^34^ GSEA revealed an enrichment of Lgr5+ stem cell signature-related genes for the factor 2 loadings (Fig. 5c) (FDR=0.002, NES=1.74) among other biological processes (Supplemental Fig. S7a). In terms of genetic mutations, ERBB2 mutation status had the strongest positive contribution to factor 2 loadings (Supplemental Fig. 6c).

**Fig. 5:**
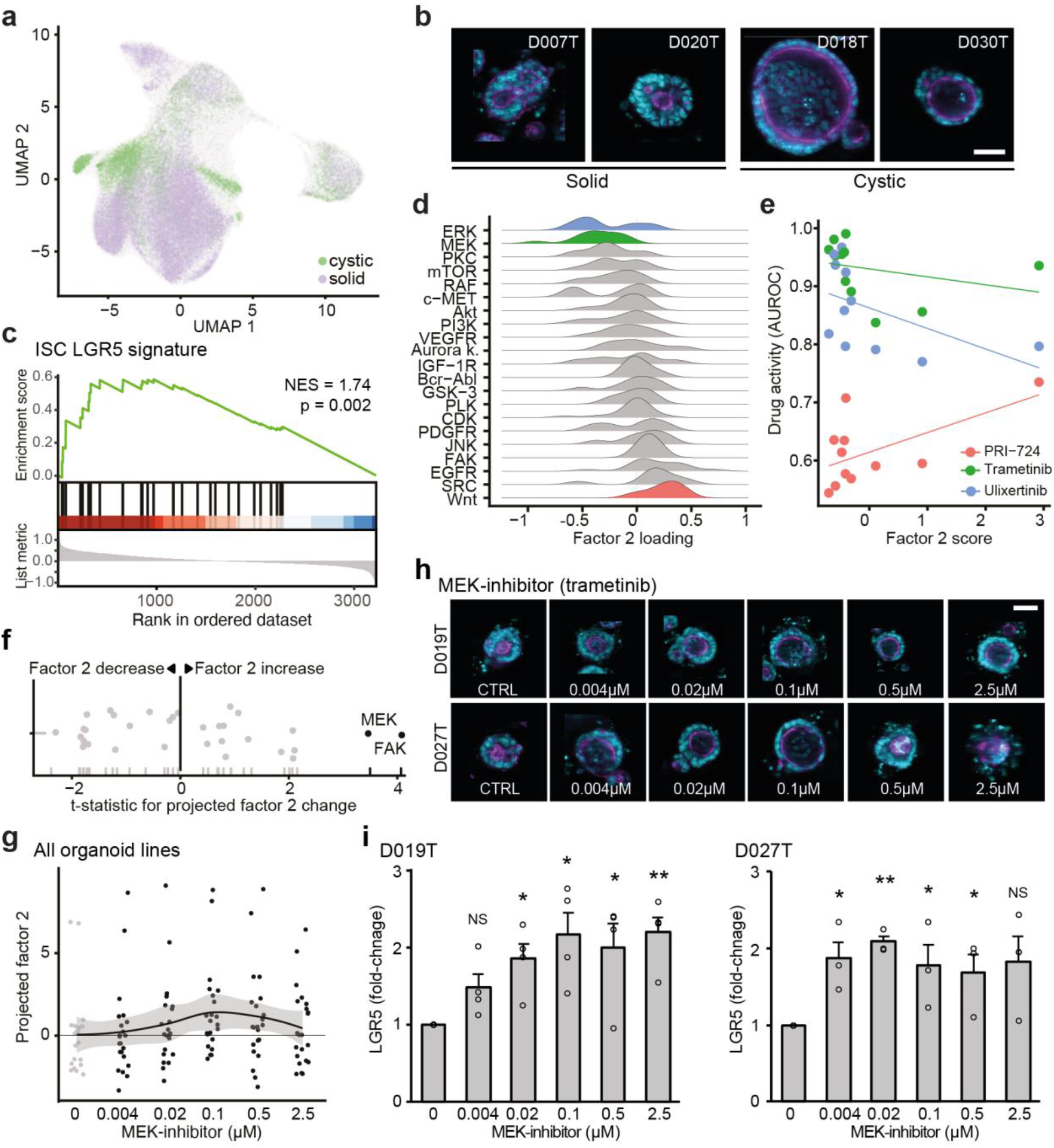
An LGR5+ stemness program is associated with cystic organoid architecture and can be induced by inhibition of MEK. **a**, UMAP visualization of cystic and solid organoid morphology in baseline state (DMSO-treated) as defined by factor 2 scores. **b**, Example images of cystic (right) and solid organoid lines. Images were automatically cropped and embedded in black background. Cyan = DNA, magenta = actin; scale bar: 50µm. **c**, Gene set enrichment analysis of the LGR5+ intestinal stem cell signature^34^ over ranked factor 2 gene expression loadings (ranking from high factor 2 loading to low factor 2 loading, NES = normalized enrichment score). **d**, Distributions of drug activity loadings for factor 2 grouped by drug targets. **e**, Relationship of representative drugs’ activity with factor 2 score. Further samples can be found in Fig. S7. **f**, Projection of factor 2 scores for drug-induced phenotypes. Highlighted are drug targets leading to a significant change in projected factor scores across all organoid lines (ANOVA). **g**, Projected dose-dependent changes in factor 2 scores after treatment with the MEK inhibitor binimetinib across organoid lines. **h**, Dose-dependent changes in organoid morphology after treatment with the MEK inhibitor trametinib. Shown are images of organoid lines D019T and D027T (cyan = DNA, magenta = actin; sampled images were cropped and embedded in black background; scale bar: 50µm) **i**, Dose-dependent changes in LGR5 transcript abundance after treatment with the MEK inhibitor trametinib, as assessed by qPCR, data from 3 (D027T) and 4 (D019T) independent replicates are presented as mean + s.e.m. * p<0.05, ** p<0.005, NS = not significant, two-sided Student’s t-test.

**Fig. 6:**
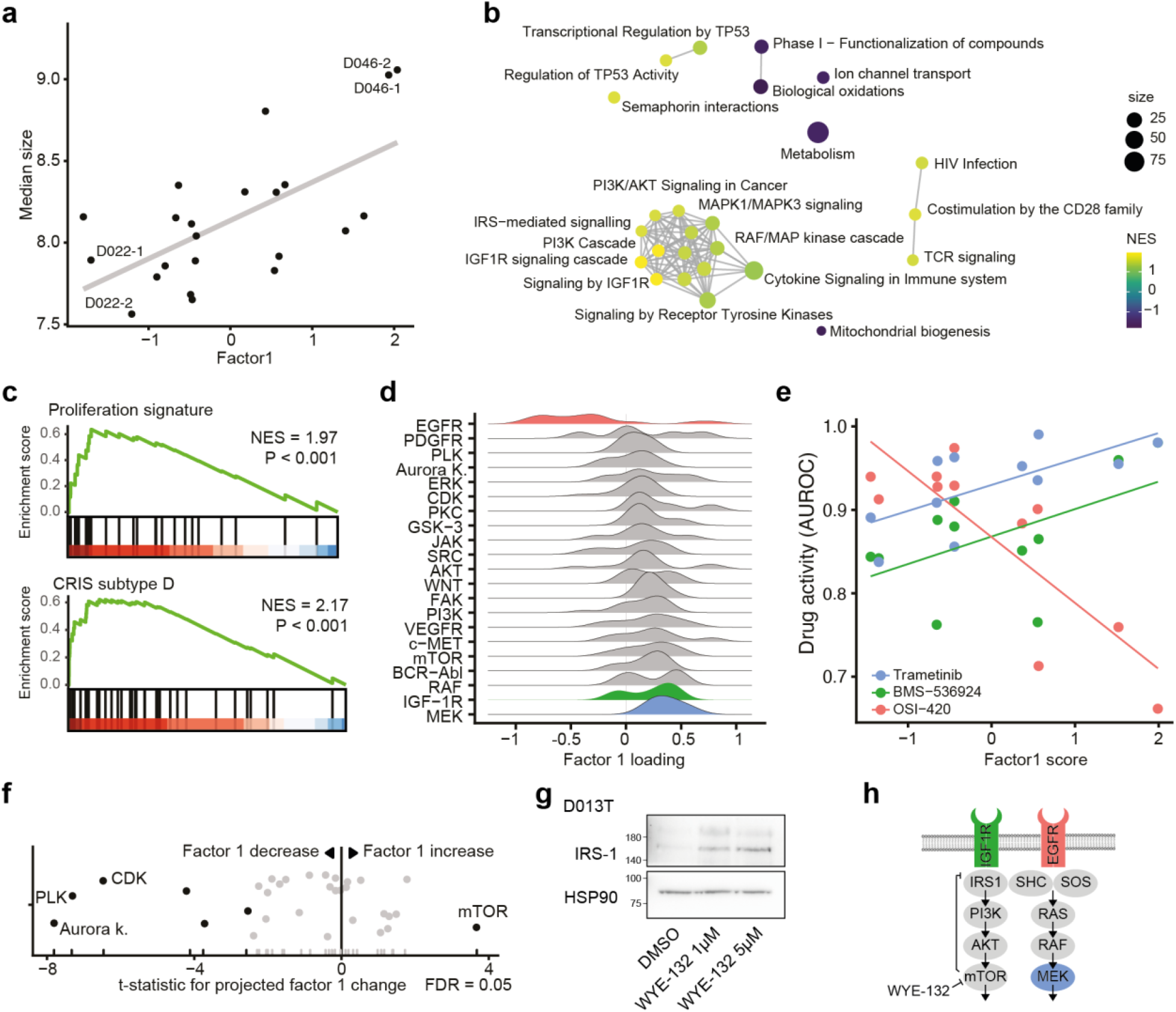
An IGF1R signaling program is associated with increased organoid size, EGFR inhibitor resistance and can be induced by mTOR inhibition. **a**, Association of factor 1 with organoid size. **b**, Gene-set enrichment network of factor 1 gene expression loadings. An edge connects Reactome pathways with more than 20% overlap. **c**, Gene set enrichment results of the “proliferation” intestinal signature^34^ and the colorectal cancer CRIS-D subtype^35^ over ranked factor 1 gene expression loadings (ranking from high factor 1 loading to low factor 1 loading, NES = normalized enrichment score). **d**, Distributions of drug activity loadings grouped by drug targets for factor 1. **e**, Relationship of selected drugs’ activity (AUROC) with factor 1 score. Further examples can be found in Fig. S8. **f**, Projection of factor 1 scores for drug-induced phenotypes. Highlighted are drug targets leading to a significant change in projected factor scores across all organoid lines (ANOVA). **g**, Western blot of IRS-1 protein abundance under ATP-competitive mTOR inhibition. A representative blot of three biological replicates with organoid line D013T is shown. **h**, Illustration of IGF1R signaling pathway with highlighted drug targets. Shown is the disinhibition of mTOR mediated IRS-1 repression by ATP competitive mTOR inhibitors.

Next, we asked if factor 2 was associated with particular drug activity or inactivity patterns. As previously described, we used the performance of a logistic regression model as drug activity score (AUROC) (Fig. 3a). Activity of Wnt signaling inhibitors and EGFR inhibitors were the strongest average contributors to a positive factor 2 score (t statistic = 3.02, FDR = 0.046 and t statistic = 3.08, FDR = 0.046, respectively), while activity of ERK and MEK inhibitors were associated with a low factor 2 score (Fig. 5d), albeit not significantly. To summarize, factor 2 high organoid lines showed an increased expression of LGR5 and were more sensitive to Wnt signaling inhibitors, such as the CBP/beta-catenin inhibitor PRI-724 (Fig. 5e and Supplemental Fig. S7b) overall suggesting increased dependency on Wnt signaling in the factor 2 high organoid state.

Prompted by the visual observation that MEK inhibitor treatment led to a related cystic architecture in organoids (Fig. 3j), we hypothesized that compound treatments could influence the plasticity between the observed organoid states. Thus, we tested whether drug treatments shifted organoid phenotype profiles in the previously defined factor space. To test for shifts in factor space, we used the previously estimated factor loading matrix for unperturbed organoid morphology, which was generated during MOFA training, as a starting point. By projecting the average phenotypic profiles of drug-treated organoids onto the factors learnt by MOFA, we were able to approximate the influence various drug treatments had on biological programs previously identified in unperturbed organoids. We observed MEK and focal adhesion kinase inhibitors significantly shifted tested organoid lines towards higher factor 2 scores (Fig. 5f and Supplemental Fig. S7c). This change in factor 2 scores was concentration dependent for MEK inhibitors (Fig. 5g and Supplemental Fig. S7d-e) and corresponded to a visual shift in organoid morphology (Fig. 5h), which was most noticeable at concentrations of 100nM (p=0.017, Fig. 5h, Supplemental Fig. S7e). Given the observation that factor 2 was enriched for an LGR5+ stem cell signature (Fig. 5c), we measured the expression of LGR5 transcripts at different concentrations of MEK inhibitor treatment for two organoid lines with representative factor scores (D019T and D027T). We observed analogous dose-dependent increases in transcript abundance (Fig. 5i). These findings were in concordance with the observation that MEK inhibitor activity had a negative contribution to factor 2 (Fig. 5d): While organoids are shifted to a factor 2 high state via MEK inhibition, within the factor 2 high state itself, organoids are relatively insensitive to this class of inhibitors. In summary, we observed an organoid state with cystic architecture, increased expression of LGR5+ stem cell related genes and increased sensitivity to Wnt signaling inhibitors that could be induced by MEK inhibition.

### An IGF1R signaling program is associated with increased organoid size, decreased EGFR inhibitor activity, and can be induced by mTOR inhibition

Next, we set out to identify the mechanisms underlying and modulating factor 1. We had previously observed that organoid size was influenced by both organoid line and drug treatments and was associated with factor 1 scores (Fig. 6a). An unsupervised gene set enrichment analysis (GSEA) for Reactome pathways across factor 1 loadings showed an enrichment for IGF1R signaling and mitogen-activated protein kinase signaling related genes. In fact, transcripts belonging to the IGF imprinting control region, H19 (rank 1) and IGF2 (rank 13), were among the strongest contributors to factor 1. This increase in proliferative signaling was confirmed by GSEA of a previously identified intestinal proliferation signature.^34^ To better understand clinical correlates to the identified gene expression patterns, we tested for molecular subtypes stemming from an analysis of cancer-cell intrinsic gene expression profiles.^35^ Factor 1 showed an enrichment for CRIS D, a molecular subtype linked to IGF2 overexpressing tumors with resistance to EGFR inhibitor therapy (Fig. 6c), and a depletion for CRIS C, which has been linked to EGFR dependency (Supplemental Fig. S8a). In fact, activity of EGFR inhibitors was the strongest contributor to a negative factor 1 score while IGF1R and MEK inhibitor activity contributed to a positive factor 1 score (Fig. 6d-e, Supplemental Fig. 8b-d). When assessing the contribution of somatic mutations, activating mutations of NRAS had the strongest contribution to a high factor 1 score (Supplemental Fig. S6c).

Next, we again used phenotype profiles of drug treated organoids and approximated how drug treatment shifted organoids along the factor 1 program. We observed a group of cell cycle related kinase inhibitors targeting polo like kinases, Aurora kinases and cyclin dependent kinases that shifted organoids to a low factor 1 score. In contrast, mTOR inhibitor treatment increased factor 1 scores in cancer organoids (Fig. 6f and Supplemental Fig. S8e). Given the observation that factor 1 was associated with IGF1R signaling and mTOR inhibitor treatment led to an increase in factor 1 scores, we hypothesized that mTOR inhibition leads to a reactive upregulation of IGF1R signaling in cancer organoids. In fact, inhibition of mTOR signaling had previously been linked to transcriptional disinhibition of IRS-1 in a negative feedback loop^36^ and reactive induction of IGF1R signaling had previously been described as a resistance mechanism to small molecule mTOR inhibitors in cancer.^37^ When testing this hypothesis in patient derived organoids, we observed a dose-dependent increase of IRS-1 protein abundance in organoids treated with the ATP competitive mTOR inhibitor WYE-132 (Fig. 6g). To summarize our findings, we observed an organoid state marked by large organoid size, elevated IGF1R dependent mitogenic signaling and relative inactivity of EGFR inhibitors. This state was inducible by inhibition of a mTOR dependent negative feedback loop in patient derived cancer organoids.

## Discussion

Organoids are currently the most complex *in vitro* cancer models with high morphological and molecular similarity to their origin and can be established from a wide variety of tumors and normal tissue.^6,7,10,14,38–40^. Given the benefits in culture efficiency and high model representativeness in comparison to conventional cell lines, it is likely that the next generation of *in vitro* cancer model cohorts are organoid based.^41,42^ Previous studies have successfully used PDOs to perform small- and medium-scale drug testing with ATP-based cell viability readouts.^7,9–15,43–45^ Additionally, imaging studies with organoids have been used to characterize developmental processes such as the self-organization of intestinal cells^25,46^ or the morphological response to individual drugs.^24,47^

While image-based profiling of *in vitro* models has become an important tool for the analysis of biological processes, particularly in drug discovery and functional genomics^17–19^, performing such high-content experiments in *ex vivo* disease models, which cannot be cultured and perturbed in 2D, has been a technological challenge. In this study, we used image-based profiling to systematically map heterogenous phenotypes of patient derived cancer organoids and their response to small-molecule perturbations. We collected data on approximately 5.5 million single organoids from 11 different colon cancer patients with >500 different small molecule perturbations. The morphology of untreated patient derived cancer organoids varied extensively within and between organoid donors. Despite the heterogeneity, organoids from different patients and perturbations showed overlapping morphological distributions, which shifted as a response to perturbation. Organoid morphology revealed compound mode-of-action and when integrated with additional biological measurements gave insight into the first set of principles governing cancer organoid architecture and plasticity. As a result, we identified two shared axes of variation for colorectal cancer organoid morphology (organoid size and cystic vs. solid architecture), their underlying biological mechanisms (IGF1R signaling and Wnt signaling), and pharmacological interventions able to move organoids along them (mTOR inhibition and MEK inhibition).

Cancer stem cells play a central role in cancer recurrence and metastasis.^48^ In colorectal cancer, cells with cancer stem cell identity are LGR5 positive.^49^ Organoid models enriched for an LGR5+ stem cell signature presented with a characteristic cystic architecture and were sensitive to inhibitors of Wnt signaling. This LGR5+ organoid state was also linked to a reduced sensitivity towards MEK inhibitors, a potential consequence of the already suppressed ERK signaling activity that has been linked to Wnt signaling in colorectal cancer.^50^ In fact, pharmacological MEK inhibition led to a shift in organoids towards a LGR5+ state, an effect that we have previously described.^33^ The use of MEK inhibitors together with Wnt signaling activating GSK3 inhibitors is an established method to maintain embryonic stem cells in vitro.^51^ A related MEK inhibitor dependent modulation of stemness in Wnt signaling dependent colon tissue may in part explain the limited success of using MEK inhibitors as monotherapy in colorectal cancer.

Insulin-like growth factors are central and conserved regulators promoting cell size, organ size and organism growth.^52,53^ The IGF1 receptor (IGF1R) signaling cascade is activated in around 20% of colorectal cancer patients and leads to downstream mitotic stimuli via mitogen activated kinase signaling and mTOR.^54^ In patient derived cancer organoids, we observed that organoid size was positively correlated with elevated IGF1R signaling activity. In accordance with clinical observations,^35^ colorectal cancer organoids in a high IGF1R signaling state were less responsive to EGFR inhibitors and more responsive to IGF1R and MEK blockade, demonstrating the central role of IGF1R mediated mitogen activated protein kinase (IGFR1-MAPK) signaling. In fact, combined blockade of MEK and IGF1R has recently been demonstrated to be a synergistic drug combination across colorectal cancer cell lines ^55^ and reciprocal resistance between IGF1R and EGFR signaling inhibitors has been described in multiple cancer types.^56^ Also, organoids could be moved into a state of increased IGF1R-MAPK signaling by inhibition of mTOR, a downstream mediator of IGF1R activity. In line with this observation, a reactive induction of IGF1R signaling has been previously described as a resistance mechanism to small molecule mTOR inhibitors in cancer.^37,57^ The emerging role of IGF1R signaling in organoid culture was recently emphasized by the observation that addition of the IGF1 ligand, relative to EGF, increased culture efficiency of organoids from healthy human small intestinal tissue.^58^

Statistical representation learning methods such as MOFA factorize a distribution of observations spanning multiple data modalities. In other words, MOFA learns factors that capture correlations between diverse biological features and scores observations along these factors. Learning factors helped identify relationships between biological processes, such as the link between organoid size, IGF1R signaling and sensitivity to IGF1R inhibitors. In search for treatments that led to drug-induced phenotype change along factors, we extended the application of factor-learning to factor-projection. This enabled us to identify mTOR and MEK inhibitors as modulators of factor 1 and factor 2, respectively. Given the fact that observations during factor-learning were sampled from a distribution of unperturbed organoids while factor-projection was done on observations from a overlapping, but distinct, distribution of perturbed organoids, our projections of perturbed organoid profiles are limited to the axes of variation defined during factor-learning. As a consequence, we are unable to observe causal relationships between factors and interventions, but only generate hypotheses based on observational data. For example, CDK inhibitor treatment reduced the score of factor 1 across all organoid models. It is, however, unlikely the reduction in factor 1 was due to CDK being an upstream regulator of IGF1R signaling. Instead CDK might serve as the dependent, mediating variable (IGFR1 signaling -> CDK signaling -> organoid size) or an independent contributor to organoid size (IGFR1 signaling-> organoid size and CDK signaling -> organoid size). Despite limitations, we believe that the approach of interpreting drug-induced phenotypes using a multi-omics representation of untreated in-vitro models is applicable to other large image-based profiling data of multiple heterogeneous *in vitro* models. This approach could potentially be further extended using causal representation learning methods that increase the understanding of cellular signaling mechanisms, the way they shape cellular morphology and how they change under various treatments during drug discovery.

While our image-based profiling study is limited by the number of studied organoid lines and organoid-level imaging resolution, our work is, to our knowledge, a first comprehensive mapping of patient derived cancer organoid morphologies across 11 organoid donors and >500 small molecule perturbations at single organoid resolution. We identified two key axes of morphological variation in cancer organoids, their underlying biological processes and pharmacological perturbations that move organoids along these axes. Previously, primary cells of monogenic diseases have been intensively studied using image based profiling for drug discovery.^59^ Our work opens up new directions for image-based profiling of complex *in vitro* disease models, as we believe this work could be expanded to search for therapeutics in somatic multigenic disease models, for example stepwise genetically edited organoid models of early colorectal cancer^31,32^, or larger cohorts of patient derived cancer organoids. In addition, more complex cellular interactions such as the interaction of the immune system with solid tumors could be explored.^60,61^ A better understanding of organoid phenotypes and the ability to use multi-omics data to annotate organoid states and their plasticity have the potential to further accelerate image-based drug discovery for complex multigenic diseases such as colorectal cancer.

## Methods

### Patients

All patients were recruited at University Hospital Mannheim, Heidelberg University, Mannheim, Germany. We included untreated patients with a new diagnosis of colon or rectal cancer in this study and obtained biopsies from their primary tumors via endoscopy. Exclusion criteria were active HIV, HBV or HCV infections. Biopsies were transported in phosphate buffered saline (PBS) on ice for subsequent organoid extraction. Clinical data, tumor characteristics and molecular tumor data were pseudonymized and collected in a database. The study was approved by the Medical Ethics Committee II of the Medical Faculty Mannheim, Heidelberg University (Reference no. 2014-633N-MA and 2016-607N-MA). All patients gave written informed consent before tumor biopsy was performed. In total, we extracted PDOs from 13 patients with colorectal cancer for this study. Patient characteristics including age, sex, tumor location, stage and treatment data can be found in Supplementary Table 1.

### Organoid culture

Organoid cultures were extracted from tumor biopsies as reported by Sato et al.^5^ with slight modifications. In short, tissue fragments were washed in DPBS (Life technologies) and digested with Liberase TH (Roche) before embedding into Matrigel (Corning) or BME R1 (Trevigen). Advanced DMEM/F12 (Life technologies) medium with Pen/Strep, Glutamax and HEPES (basal medium) was supplemented with 100 ng/ml Noggin (Peprotech), 1x B27 (Life technologies), 1,25 mM n-Acetyl Cysteine (Sigma), 10 mM Nicotinamide (Sigma), 50 ng/ml human EGF (Peprotech), 10 nM Gastrin (Peprotech), 500 nM A83-01 (Biocat), 10 nM Prostaglandin E2 (Santa Cruz Biotechnology), 10 µM Y-27632 (Selleck chemicals) and 100 mg/ml Primocin (Invivogen). Initially, cells were kept in 4 conditions including medium as described (ENA), or supplemented with additional 3 uM SB202190 (Biomol) (ENAS), 50% Wnt-conditioned medium and 20% R-Spondin conditioned medium (WENRA) or both (WENRAS), as described by Fujii et al.^6^ The tumor niche was determined after 7-14 days and cells were subsequently cultured in the condition with best visible growth. PDOs were passaged every 7-10 days and medium was refreshed every 2-3 days. 13 PDO lines were analyzed within this study, data of all PDO lines including niche and growth rate are denoted in Supplementary Table 1.

### Amplicon sequencing

DNA was isolated with the DNA blood and tissue kit (Qiagen). Sequencing libraries were prepared with a custom panel (Tru-Seq custom library kit, Illumina) according to the manufacturer’s protocol and sequenced on a MiSeq (Illumina) as reported previously.^62^ Targeted regions included the most commonly mutated hot spots in colorectal cancer in 46 genes captured with 157 amplicons of approximately 250bp length. After mapping the reads to the GRC38 reference genome using Burrows-Wheeler Aligner (BWA), data were analyzed using the Genome Analysis Toolkit (GATK).^63^ Base recalibration was performed and variants were called using the MuTect2 pipeline. Variants with a variant frequency below 10%, with less than 10 reads, or with a high strand bias (FS<60) were filtered out. Variants were annotated with Ensembl variant effect predictor^64^ and manually checked and curated using integrative genomics viewer, if necessary.^65^ Only non-synonymous variants present in COSMIC^66^ were considered true somatic cancer mutations. Also, all variants annotated “benign” according to PolyPhen database and “tolerated” in SIFT database were excluded, as well as variants with a high frequency in the general population as determined by a GnomAD^67^ frequency of >0.001.

### Expression profiling

Organoid RNA was isolated with the RNeasy mini kit after snap freezing organoids on dry ice. Samples were hybridized on Affymetrix U133 plus 2.0 arrays. Raw microarray data were normalized using the robust multi-array average (RMA) method^68^ followed by quantile normalization as implemented in the ‘affy’^69^ R/Bioconductor package. In order to exclude the presence of batch effects in the data, principal component analysis and hierarchical clustering were applied. Consensus molecular subtypes were determined as described previously^70^ using the single sample CMS classification algorithm with default parameters as implemented in the R package ‘CMSclassifier’. In all cases, differential gene expression analyses were performed using a moderated t-test as implemented in the R/Bioconductor package ‘limma’.^71^ Gene set enrichment analyses were performed using ConsensusPathDB^72^ for discrete gene sets or GSEA as implemented in the ‘fgsea‘^73,74^ R/Bioconductor package for ranked gene lists. Wikipathways^75^ or Reactome^76^ were used for pathway analysis. Gene expression analysis was done in R version 4.0.0. When possible, packages were installed via bioconductor.

### Compound profiling

#### Cell seeding

PDO drug profiling followed a standardized protocol with comprehensive documentation of all procedures. Organoids were collected and digested in TrypLE Express (Life technologies). Fragments were collected in basal medium with 300 U/ml DNAse (Sigma) and strained through a 40µm filter to achieve a homogeneous cell suspension with single cells and small clusters of cells, but without large organoid fragments. 384 well µclear assay plates (Greiner) were coated with 10µL BME V2 (Trevigen) at a concentration of 6.3 mg/ml in basal medium, centrifuged and incubated for >20 min at 37° C to allow solidification of the gel. PDO cell clusters together with culture medium (ENA) and 0,8 mg/ml BME V2 were added in a volume of 50µl per well using a Multidrop dispenser (Thermo Fisher Scientific). Plates were sealed with a plate-loc (Agilent) and centrifuged for an additional 20 min allowing cells to settle on the pre-dispensed gel. Cell number was normalized before seeding by measuring ATP levels in a 1:2 dilution series of digested organoids with CellTiter-Glo (Promega). The number of cells matching 10,000 photons (Berthold Technologies) was seeded in each well. After seeding of organoid fragments, plates were incubated for three days at 37°C to allow organoid formation before addition of small molecules. Two biological replicates (defined as an independent passage) of each PDO line were profiled. Mean passage number of the PDO lines by the time of profiling of the first replicate was 9 (median 9) and PDOs were passaged up to two more times before the second replicate. In total, 13 PDO lines underwent profiling with the clinical cancer library and the KiStem library with high throughput imaging. Data from two organoid lines (D015T, D021T) later had to be excluded due to too many out-of-focus organoids (more details below). One line, D020T, was profiled twice within different experimental batches (D020T01 and D020T02). If not shown otherwise, data from D020T01 was used.

#### Compound libraries

Two compound libraries were used for screening: A library containing 63 clinically relevant small molecules (clinical cancer library, Supplementary table 3) and a library of 464 compounds targeting kinases and stem cell or developmental pathways associated genes (Ki-Stem library, Supplementary table 4). The clinical cancer library was manually curated by relevance for current (colorectal) cancer therapy, mechanism of action and potential clinical applicability. small molecules of this library were mainly in clinical use or in phase I/II clinical trials. Five concentrations per compound were screened (five-fold dilutions). The concentrations were determined by analysis of literature data from previous 3D and 2D drug screens and own experiments. All small molecules within the KiStem library were used in a concentration of 7.5µM. All small molecules were obtained from Selleck chemicals. Libraries were arranged in an optimized random layout. We stored compound libraries in DMSO at -80 C.

#### Compound treatment

30µl medium was aspirated from all screening plates and replaced with fresh ENA medium devoid of Y-27632, resulting in 45µl volume per well. Drug libraries were diluted in basal medium and subsequently 5µl of each small molecule was distributed to screening plates. All liquid handling steps were performed using a Biomek FX robotic system (Beckmann Coulter). Plates were sealed and incubated with small molecules for four days.

#### Luminescence viability read out

Plates undergoing viability screening were treated with 30µl CellTiter-Glo reagent after medium aspiration with a Biomek FX (Beckmann Coulter). After incubation for 30 minutes, luminescence levels were measured with a Mithras reader (Berthold technologies).

#### Image-based phenotyping

Image-IT DeadGreen (Thermo Fisher) was added to the cultures with a Multidrop dispenser (Thermo Fisher) in 100nM final concentration and incubated for 4 hours. Afterwards, medium was removed, and organoid cultures were fixed with 3% PFA in PBS with 1% BSA. Fixed plates were stored at 4° C for up to 3 days before permeabilization and staining. On the day of imaging, organoids were permeabilized with 0.3% Triton-X-100 and 0.05% Tween in PBS with 1% BSA and stained with 0.1µg/ml TRITC-Phalloidin (Sigma) and 2µg/ml DAPI (Sigma). All liquid handling steps were performed with a BiomekFX (Beckmann Coulter). Screening plates were imaged with an Incell Analyzer 6000 (GE Healthcare) line-scanning confocal fluorescent microscope. We acquired 4 fields per well with z-stacks of 16 slices at 10x magnification. The z-steps between the 16 slices had a distance of 5µm, the depth of field of each slice was 3.9µm.

### Immunohistochemistry

PDOs were fixed for 20 min in 4% (v/v) Roti Histofix (Carl Roth) followed by embedding into MicroTissues 3D Petri Dish micromolds (Sigma Aldrich) using 2 % (w/v) Agarose LE (Sigma) in PBS supplemented with 0.5 mM DTT. Thereafter, PDOs were subjected to dehydration steps and embedding in paraffin. Formalin-fixed agarose/paraffin-embedded sections (3-5µm) were manually cut from blocks with a microtome (Leica RM 2145) and transferred to glass slides (Superfrost, Thermo Fisher Scientific) before H&E staining using automated staining devices.

### Realtime quantitative PCR

Total RNA was isolated from organoids with the RNeasy Mini kit (Qiagen). cDNA synthesis was done with Verso cDNA kit (Thermo Fisher Scientific), and RT-PCR was performed using the SYBR Green Mix (Roche, Nutley, NJ, USA) on LightCycler480 system (Roche). The following primers for *LGR5* were used: 5’-TTC CCA GGG AGT GGA TTC TAT-3’ (forward) and 5’-ACC AGA CTA TGC CTT TGG AAA C-3’ (reverse). Results were normalized to *UBC* mRNA using 5’-CTG ATC AGC AGA GGT TGA TCT TT-3’ forward and 5’-TCT GGA TGT TGT AGT CAG ACA GG-3’ reverse primers.

### Western Blot

Organoids seeded in 6-well plates were harvested after 3-days incubation with WYE-132 in RIPA buffer (Thermo Scientific) supplemented with protease inhibitors (Complete Mini, Roche) and phosphatase inhibitors (Phosphatase Inhibitor 1 and 2, Sigma), followed by sonication (Branson Sonifier, Heinemann). Protein concentrations of supernatants were measured using a BCA assay kit (ThermoFischer Scientific). Lysates were mixed with an SDS-loading buffer and heated to 99°C for 5 minutes. Proteins were separated by SDS–PAGE in MOPS running buffer and transferred to a nitrocellulose membrane. Membranes were blocked with 5% (w/v) skim milk in PBS containing 0.1% (v/v) Triton X-100 (PBS-T). Antibodies against IRS1 (06-248, Sigma-Aldrich) and HSP-90 (sc-13119, Santa Cruz) as loading control were used in 1:1000 dilution in 5% milk in PBS-T, secondary antibodies (Mouse IgG HRP ECL, Sigma Aldrich) were used in 1:10000. ECL Western Blotting W1001 (Promega) was used for visualization of bands.

### Image analysis

#### Image processing

Microscopic image z-stacks were illumination corrected using a prospective method, compressed to HDF5 format and underwent maximum contrast projection using the *MaxContrastProjection* package for further processing of the images. This algorithm projects the multi-channel 3D image stack onto a plane by retaining the pixel information with the strongest contrast to its neighboring pixels. We used a two-step procedure to establish segmentation: First, organoids were segmented using a model based on fluorescence channel intensity. The intensity segmentati on was then used to perform weakly supervised learning with a deep convolutional neural network (CNN) for object identification on the partially correct intensity segmentation, leveraging the robustness of CNNs with regard to mislabeled training data and eliminating the need for expensive manual annotations. For further analysis, we used a model-free outlier detection to remove segmented objects with a particle size of 300 pixels and lower to remove non-organoid objects. Standard image features, including shape, moment, intensity, and Haralick texture features ^77^ on multiple scales, were extracted using the R/Bioconductor package *EBImage*.^*78*^. Initially, we extracted a total of 1572 features for each individual organoid image. However, texture features were meaningless for scales larger than the actual organoid size. To simplify the analysis, we thus only retained texture features that were well defined for all organoids and on a scale smaller than the smallest organoids in the image dataset. This ensured that the dataset contained no NA-values requiring imputation. A feature was considered “well-defined’ if the median absolute deviation across the entire dataset was strictly greater than 0. In other words, if more than half of all organoids exhibited an identical value for a feature, then that feature was discarded for further analysis. This resulted in 528 well-defined features. We did not perform feature selection based on between-replicate correlation of well-averaged features as we used single-organoid features for further analysis and selected downstream methods used (Random-Forest) did not require pre-selection of features or were based on principal components across the complete dataset (logistic regression). To allow comparison between various PDO lines and drug perturbations, the distributions of features describing organoids from different batches were adjusted by centering. Out-of-focus objects were programmatically removed from the dataset using a feature based random forest classifier. Data from two PDO lines (D015T, D021T) had to be excluded from image analysis due to too many out of focus objects, resulting in 11 analyzed PDO lines. In addition, data from three individual plates (D027T01P906L03, D020T01P906L03, D013T01P001L02) were excluded from further analysis due to out-of-focus artefacts. Images were processed with R3.6.0 and packages were downloaded from bioconducotr.

#### Analysis of unperturbed organoid phenotypes

Principal components were calculated for the entire dataset using incremental principal component analysis. A set of 25 principal components were selected, explaining approx. 81% of the total variance within the dataset. Next we embedded the first 25 principal components using uniform manifold approximation and projection (UMAP) with min_distance of 0.1, 15 nearest neighbors and otherwise default monocle3 parameters.^27^ Embedded objects were clustered using the leiden graph based clustering algorithm with a resolution parameter of 10E-7.^79^ For the illustration of dose-dependent changes in organoid morphology we fitted principal curves through downsampled UMAP observations using the princurve R package.^80^

#### Live-dead classification

A random forest classifier (scikit-learn v1.0) with 10 trees was trained on the original 1572 single organoid features to differentiate living from dead organoids. Organoids treated with DMSO were used as negative (i.e. living) controls while organoids treated with Bortezomib and SN-38 at the two highest concentrations were used as positive (i.e. dead) controls. Visual inspection of the projected images confirmed our choice of positive controls. Models were trained and validated using only observations from the clinical cancer panel with a 60-40 train-validation split. Initial classification performance metrics were estimated using the validation dataset. A separate classifier was trained for each individual line to ensure inter-line independence, however individual classifiers were evaluated on validation data from foreign organoid lines to assess generalizability. Classifiers relying on less information (i.e. a combination of actin/TRITC or DNA/DAPI staining alone, compared to all three fluorescence channels) were tested by masking of input features. Binary classification results were averaged within wells to obtain viability scores ranging from 0 to 1, indicating how lethal a treatment was. This procedure was applied to the complete imaging data.

#### Analysis of drug activity and drug-induced phenotypes

A logistic regression model (scikit-learn) was trained per line and treatment (and per concentration where applicable) to differentiate treated organoids from negative controls based on the PCA-transformed features.^81^ For model training, organoid observations were separated into training and validation data with a 50-50 split. A hyperparameter grid search for L2 regularization strength was performed on the training set using 5-fold cross validation. Selected models were then trained on the validation set and model performance, expressed in the area under the receiver operating characteristic curve (AUROC), was estimated using 10-fold cross validation. Next, we selected active compound treatments in which robust morphological changes were observed in at least one line. Treatments were categorized as either active or inactive based on the performance of the logistic regression classifier. We defined a compound treatment as “active” when treated and untreated organoids in the validation dataset could be correctly identified by their corresponding classifier with an average area under the receiver operating characteristic curve (AUROC) of 0.85 or greater. The model coefficients, which can be understood as the direction of the normal vector perpendicular to the separating hyperplane in organoid feature space, was interpreted as the drug-induced effect. We chose this approach to account for the high intra- and inter-sample heterogeneity of primary patient derived organoids. We accepted the strong reference to DMSO treated organoids to describe compound treatments. Drugs were clustered based on the cosine similarity. We compared this approach to a model-free Pearson correlation-based clustering. We then aggregated compound induced phenotypic profiles across all PDO lines and applied contingency testing.^82^ Fisher’s exact test was used to identify enrichments of compounds with the same mode-of-action.

#### Analysis of dose-response relationships for organoid viability measurements

Cell Titer Glo raw data of each plate were first normalized using the Loess-fit method^83^ in order to correct for edge effects. Subsequently, each plate was normalized by division with the median viability score of the DMSO controls. For drugs tested in multiple concentrations, drug response Hill curves (DRC) were fitted and area under the curve values were calculated for each DRC using the ‘PharmacoGx’ R/Bioconductor package.^84^ The same method was used for predictions by the Live-dead classifier in cases where multiple concentrations were available.

### Multi-omics factor analysis

#### Model training

A multi-omics factor analysis model was trained based on a set of five modalities describing unperturbed organoid lines:

- organoid size estimated based on log-normal model fit of all DMSO treated organoids [22 replicates, 1 feature]
- organoid somatic mutations as determined by amplicon sequencing [20 replicates, 12 features]
- organoid gene expression including the top 10% genes with the highest coefficient of variance after robust multi-array average normalization [22 replicates, 3222 features]
- organoid morphology as determined by averaging DMSO treated morphological profiles [22 replicates, 25 features]
- organoid drug activity as determined by AUROC score of logistic regression models for drugs that were defined as active in at least one observation [22 replicates, 252 features]

Input data was scaled and the MOFA model was trained with default MOFA2 training parameters and a number of 3 factors.^30^ The number of factors was chosen given the limited number of observations in our training data. The further analysis focused on the first two factors, which correlated with prominent visible organoid phenotypes. Gene set enrichment analysis and Reactome pathway enrichment of factor loadings was performed using the clusterprofiler R package (v4.2).^85^ Enrichment of drug targets within factor loadings was tested using ANOVA by fitting a linear model, lm(factor loading ∼ target). Drug targets that were represented with at least three small molecule inhibitors were included in this analysis. The analysis was run using the MOFA docker container available from https://hub.docker.com/r/gtca/mofa2.

#### Model projection

To estimate the factor scores for drug-induced organoid morphologies, the morphology profiles of organoids treated with the same drug were averaged. The resulting average profile matrix was multipled with the pseudoinverse of the previously learnt model loading matrix for organoid morphology data. The resulting projected factor score matrix was used to estimate the drug-induced biological changes in cancer organoids. Associations between drug targets and projected factor scores of drug treated organoids were identified via ANOVA by fitting a linear model, lm(projected factor score ∼ target). Drug targets that were represented with at least three small molecule inhibitors were included in this analysis.

### Software and data availability

Software for organoid image analysis (including segmentation, feature extraction, analysis of drug-induced phenotypes, live-dead-classification), the scripts for analysis of luminescence data, dose response relationships, expression data and multi-omics factor analysis, as well as a sample of the data are available at: https://github.com/boutroslab/Supp_BetgeRindtorff_2021. Microarray data are made available in Gene Expression Omnibus (GEO, https://www.ncbi.nlm.nih.gov/geo/) under accession no. GSE117548. Amplicon sequencing data are made available through controlled access in the European Genome Phenome Archive (EGA, https://www.ebi.ac.uk/ega/home, accession no. EGAD00001004313). Data access requests for sequence data will be evaluated and transferred upon completion of a data transfer agreement and authorization by the data access committee of DKFZ and Department of Medicine II, Medical Faculty Mannheim.

## Acknowledgements

We thank all patients participating in this study as well as the teams approaching patients for consent. We thank A. Falzone, A. Kerner and K. Kaiser for excellent technical assistance, B. Velten, K. Boonekamp, C. Scheeder and F. Heigwer for helpful discussions on the manuscript and C. Cai for help with immunohistochemical stainings. We are grateful to H. Farin for helpful discussions. We are grateful to J. Hodzic, B. Huber, M. Hirth, G. Kähler, M. Sold and the Central Endoscopy Unit (ZIE) of the University Hospital Mannheim for inclusion of patients and asservation of biopsies. We thank the DKFZ Genomics and Proteomics Core Facility and the Center for Medical Research (ZMF) Mannheim for assistance with Microarrays and H/E stainings, respectively. This work was supported by the Hector Stiftung II, Germany. J.B. was supported by the “Translational Physician Scientist (TraPS)” program of the Medical Faculty Mannheim, Heidelberg University, by the Dr. Hans und Lore Graf Stiftung, Germany and by the Hector II Foundation, Weinheim, Germany. MPE is supported by the DFG (GRK2727) and the Land Baden-Württemberg (BW-ZDFP). Research in the lab of MB was in part supported by an ERC Advanced Grant and the German Network for Bioinformatics Infrastructure (de.NBI).

## Author contributions

Conceptualization, J.B., N.R., J.S., M.E. and M.B; Methodology, J.B., N.R., H.G., T.M., and F.H.; Formal Analysis, N.R., J.S., B.R., E.V.; Software, J.S., N.R. and B.F.; Investigation, J.B., N.R., C.D., H.G., F.H., K.S.-M., V.H., T.Gu., L.F., S.B., T.Ga., I.B., R.J, N.H. and T.Z.; Writing – Original Draft, J.B. and N.R.; Writing – Review & Editing, J.B., N.R., M.E. and M.B.; Data curation, J.B., N.R., J.S. and B.R.; Visualization, J.B., N.R., J.S. and B.R.; Funding Acquisition, M.E., M.B. and J.B.; Resources, M.E. and M.B.; Supervision, J.B., K.B-H., E.B., M.E. and M.B.

