## Supplemental Figures for "The drug-induced phenotypic landscape of colorectal cancer organoids"

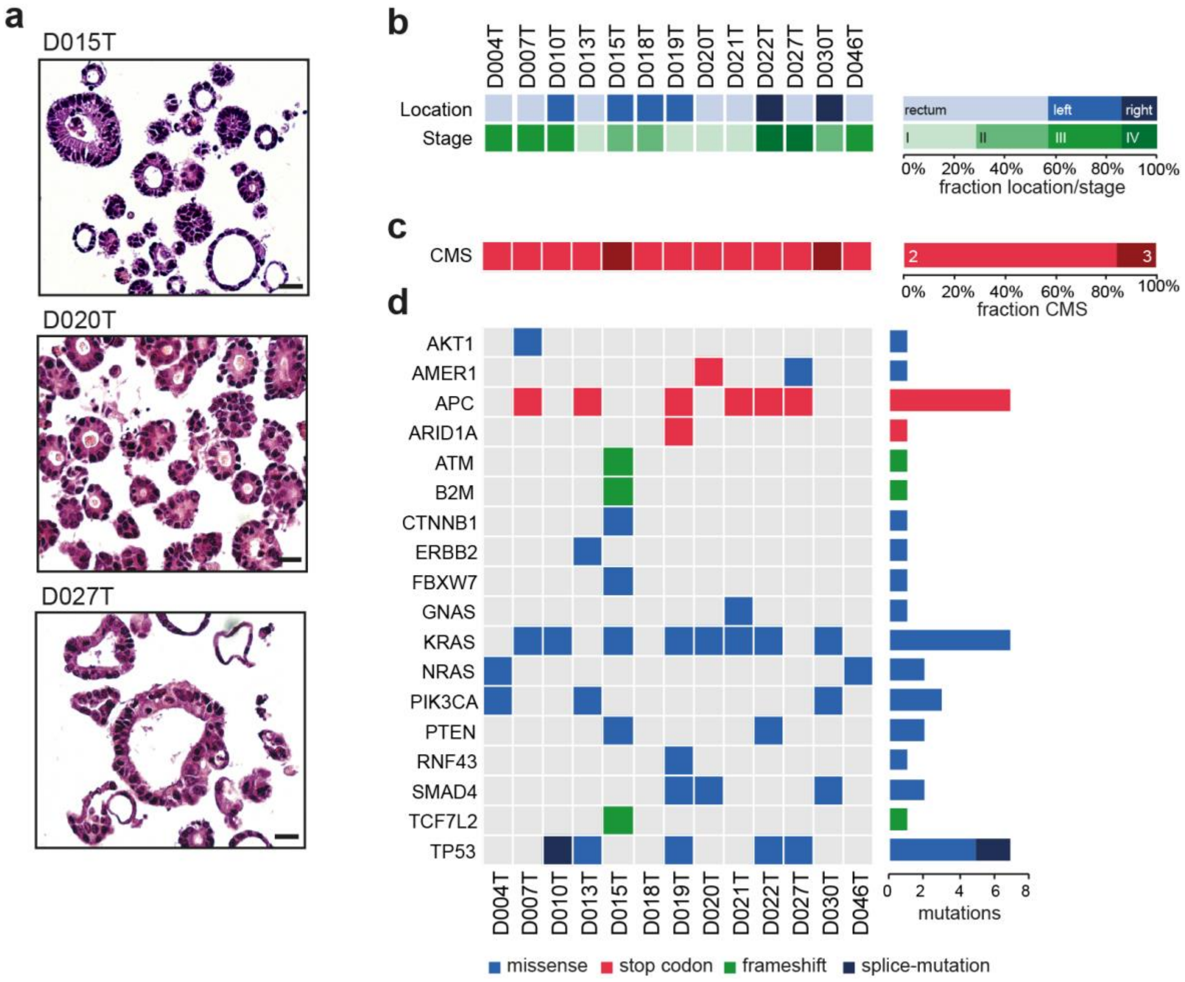


**Supplemental Fig. S1: Establishment of patient derived organoids for high-throughput image-based compound profiling**. **a**, Examples of H&E stained slides of selected representative PDO cultures; scale bar: 25µm. **b**, Tumor location (right/left/rectum) and AJCC/UICC stage of colorectal cancers that PDOs were derived from. **c**, Consensus molecular subtypes of PDOs determined by RNA expression analysis. **d**, Mutation status in PDOs, as analyzed by amplicon sequencing (more information in Supplemental Table S1).

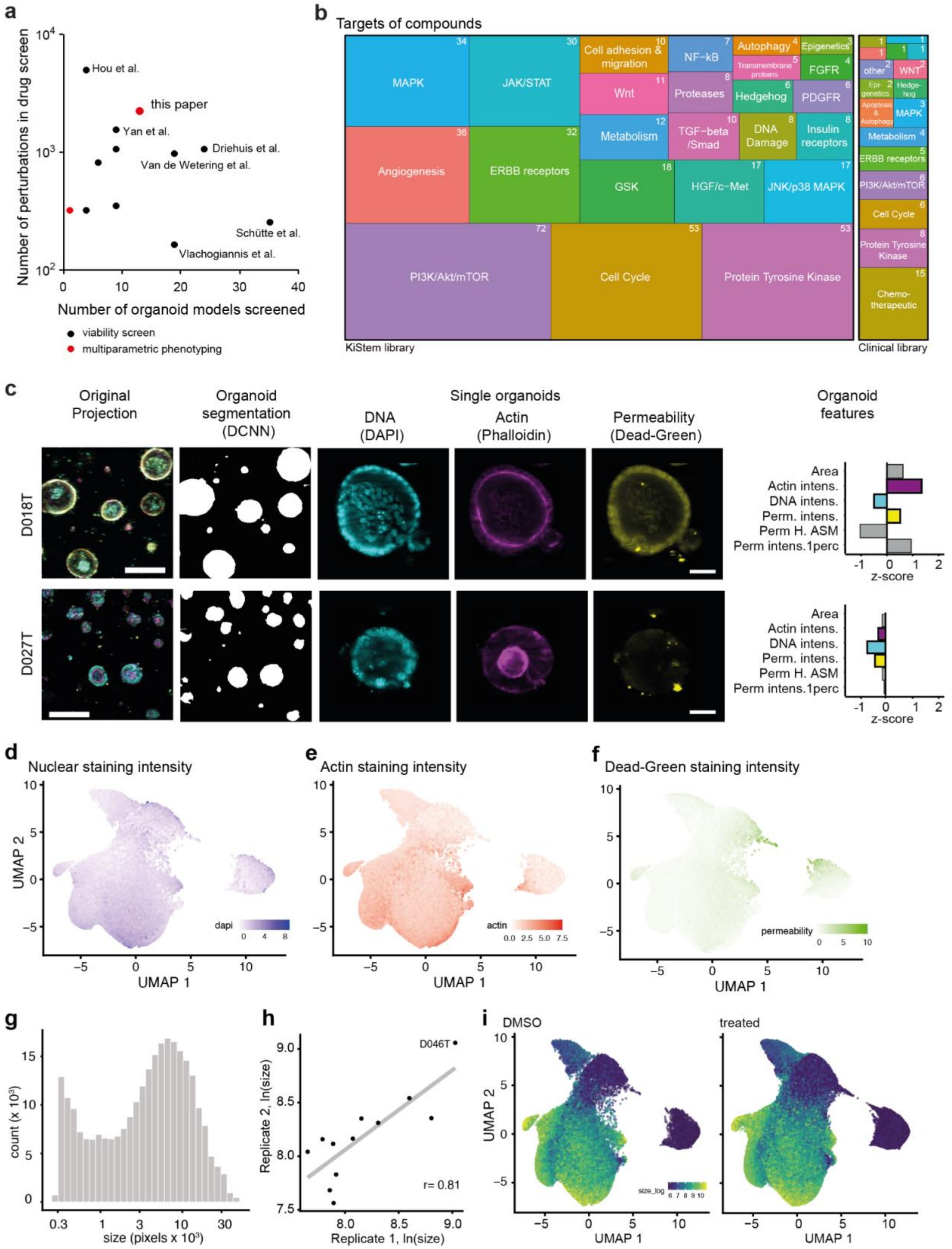

a
Number of perturbations in drug screen
Number of organoid models screened
Hou et al.
this paper
Yan et al.
Driehuis et al.
Van de Wetering et al.
Schütte et al.
Vlachogiannis et al.
viability screen
multiparametric phenotyping
b
Targets of compounds
MAPK
JAK/STAT
Cell adhesion & migration
NF-kB
Autophagy
Epigenetics
Transmembrane proteins
FGFR
Wnt
Proteases
Hedgehog
PDGFR
Angiogenesis
ERBB receptors
Metabolism
TGF-beta /Smad
DNA Damage
Insulin receptors
GSK
HGF/c-Met
JNK/p38 MAPK
PI3K/Akt/mTOR
Cell Cycle
Protein Tyrosine Kinase
KiStem library
Clinical library
other
WNT
Epi-genetics
Hedge-hog
Apoptosis & Autophagy
Chemo-therapeutic
c
Original Projection
Organoid segmentation (DCNN)
Single organoids
DNA (DAPI)
Actin (Phalloidin)
Permeability (Dead-Green)
Organoid features
D018T
D027T
Area
Actin intens.
DNA intens.
Perm. intens.
Perm H. ASM
Perm intens.1perc
z-score
d
Nuclear staining intensity
e
Actin staining intensity
f
Dead-Green staining intensity
UMAP 1
UMAP 2
dapi
actin
permeability
g
count (x 10$^3$)
size (pixels x 10$^3$)
h
Replicate 1, ln(size)
Replicate 2, ln(size)
D046T
r= 0.81
i
DMSO
treated
size_log

**Supplemental Fig. S2: Automated morphological analysis of patient derived organoids. a**, Number of organoid models and number of perturbations in previous publications reporting high-throughput drug screenings with patient derived cancer organoids **b**, Graphical representation of the compound libraries used for drug screening in this project: A library targeting kinases and stem cell pathways (KiStem library, 464 compounds) and a clinical library with 63 drugs in 5 concentrations. **c**, The image-processing pipeline illustrated with representative example images from 2 organoid lines: The multi-channel (DNA/DAPI, actin/TRITC, permeability/FITC) 3D image stack was projected onto a plane and a deep convolutional neural network subsequently recognized complete foreground organoids. Descriptive features were extracted from all three channels to quantify phenotypes. Feature plots show the median phenotype of unperturbed organoids, six example features (Area, Phalloidin intensity, DAPI intensity, FITC intensity, FITC Haralic angular second moment (ASM) and FITC intensity 1-percentile) and their z-scores relative to all profiled organoid lines are shown. **d-f**, Uniform Manifold Approximation and Projection (UMAP) of organoid-level features marked by DNA (DAPI) staining intensity (d), actin (Phalloidi/FITC) staining intensity (e) and permeability (DeadGreen) staining intensity (f). **g**, Distribution of organoid size in control (DMSO) treated organoid lines. **h**, Replicate correlation of organoid size in control treated organoids. **i**, UMAP representation of DMSO treated and drug treated organoids.

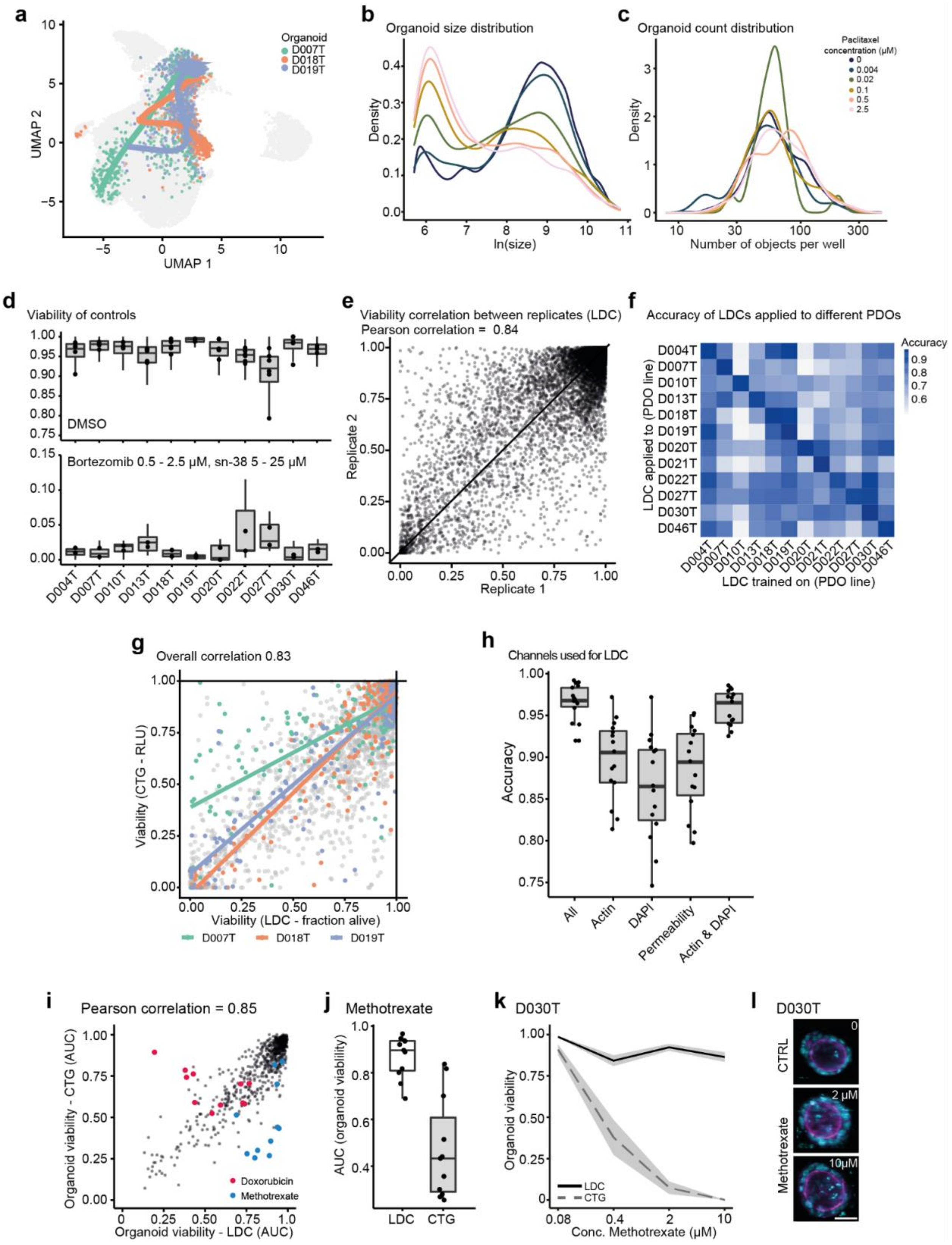
a
Organoid
D007T
D018T
D019T
UMAP 2
UMAP 1
b
Organoid size distribution
Density
ln(size)
c
Organoid count distribution
Paclitaxel concentration (μM)
0
0.004
0.02
0.1
0.5
2.5
Density
Number of objects per well
d
Viability of controls
DMSO
Bortezomib 0.5 - 2.5 μM, sn-38 5 - 25 μM
D004T
D007T
D010T
D013T
D018T
D019T
D020T
D022T
D027T
D030T
D046T
e
Viability correlation between replicates (LDC)
Pearson correlation = 0.84
Replicate 2
Replicate 1
f
Accuracy of LDCs applied to different PDOs
Accuracy
LDC applied to (PDO line)
LDC trained on (PDO line)
D004T
D007T
D010T
D013T
D018T
D019T
D020T
D021T
D022T
D027T
D030T
D046T
g
Overall correlation 0.83
Viability (CTG - RLU)
Viability (LDC - fraction alive)
D007T
D018T
D019T
h
Channels used for LDC
Accuracy
All
Actin
DAPI
Permeability
Actin & DAPI
i
Pearson correlation = 0.85
Organoid viability - CTG (AUC)
Organoid viability - LDC (AUC)
Doxorubicin
Methotrexate
j
Methotrexate
AUC (organoid viability)
LDC
CTG
k
D030T
Organoid viability
LDC
CTG
Conc. Methotrexate (μM)
l
D030T
CTRL
Methotrexate
0
2 μM
10μM

**Supplemental Fig. S3: Analysis of lethal and non-lethal PDO phenotypes. a**, Dose-dependent-trajectory of sn-38 drug effect. UMAP of organoid-level features are shown for example organoid lines D007T, D018T, D019T. For visual purposes, trajectory inference was limited to partition 1, the left-hand set of measurements within the UMAP representing ca. 95% of all imaging data. **b**, Distribution of organoid size at different concentrations of paclitaxel. Shown is a random sample of 30% of all paclitaxel treated organoids for this and the following figures. **c**, Distribution of organoid number per well at different concentrations of paclitaxel. **d-f**, Supervised machine learning of organoid viability. Organoid phenotype based viability classifiers (live-dead classifiers, LDC) were trained on positive- (high-dose bortezomib and SN-38) and negative (DMSO) controls. **d**, Fraction of DMSO treated (correctly) classified as viable (top) and fraction of organoids classified as dead in positive controls (bottom) for each PDO line. **e**, Correlation of viability (fraction of viable organoids per well) between 2 biological replicates classified by LDC. **f**, Transfer learning of organoid viability. LDCs were trained on the feature sets of negative- and positive controls for every PDO line. Receiver operator characteristic curves (ROCs) were analyzed on validation sets of negative- and positive controls of the same lines and other lines. Systematic analysis of the transfer-performance of all LDCs when applied across data from all organoid lines is shown. Classification performance is measured as the AUROC (area under the receiver operating characteristic curve). **g**, Association of organoid viability of selected example lines as determined by LDC vs. luminescence-based, ATP-dependent viability profiling with CellTiter-Glo (CTG). **h**, Accuracy of LDCs trained on image-features of all three available fluorescence channels compared to classifiers trained on single-channel data (actin/TRIC, DNA/DAPI, cell permeability/FITC) and on a combination of actin (TRIC) and DNA (DAPI) data only. **i**, Pearson correlation of areas under the dose-response curve (AUCs) for drugs with multiple tested concentrations. Shown are AUCs for phenotype based viability classification (x-axis) and CTG ground truth (r = 0.87). Measurements of methotrexate and doxorubicin are marked in blue and red, respectively. All profiled lines are included. **j**, Methotrexate, an example of a drug that had divergent viability response results between phenotype based and CTG measurements. **k**, Representative example of dose-response curves (organoid line D030T) for methotrexate. The range of values from two biological replicates are shaded in grey. **l**, Representative images of organoids treated with DMSO (control) and methotrexate 2 and 10 µM.; cyan = DAPI, magenta = Phalloidin, yellow = cell permeability; scale-bar: 200µm.

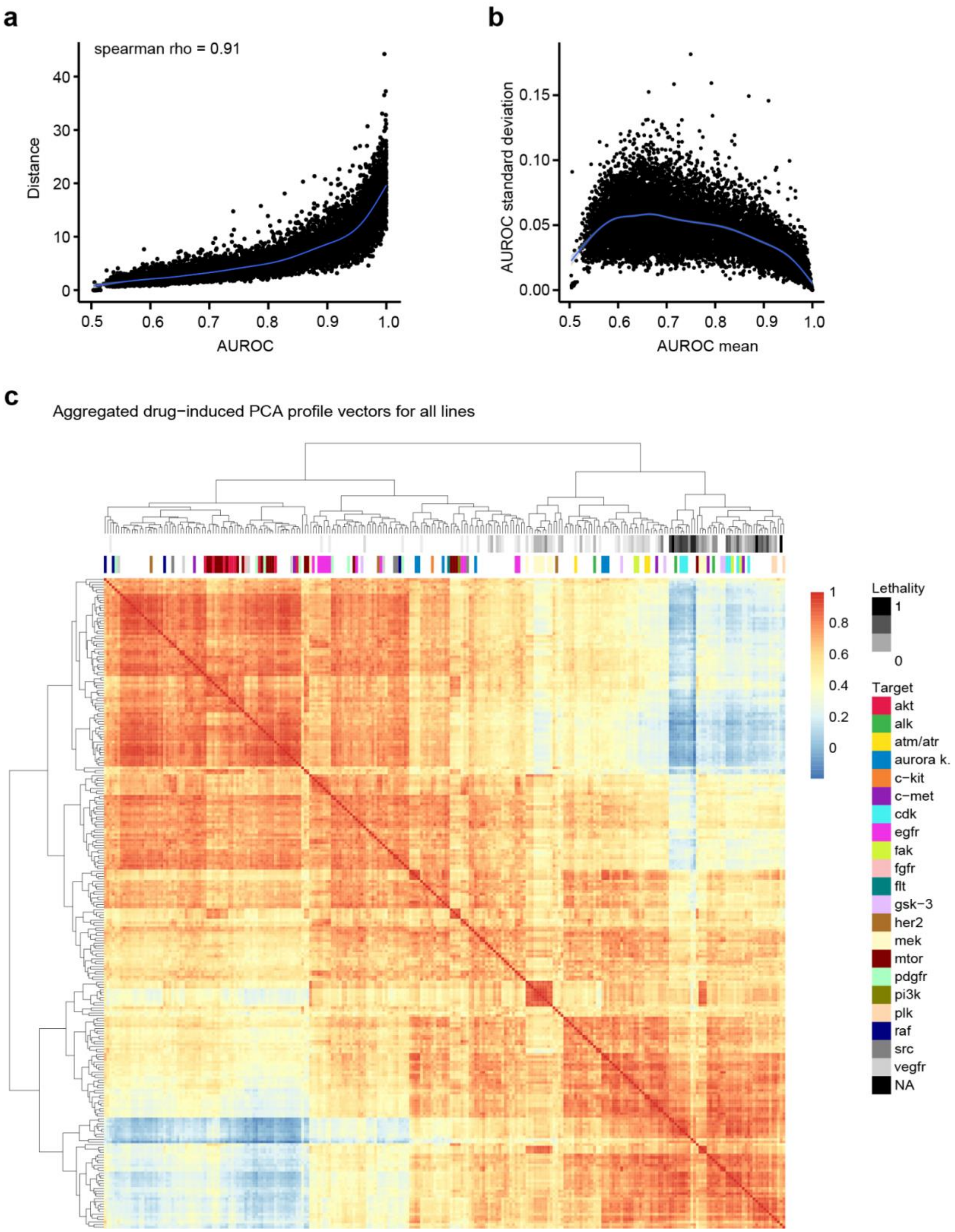


**Supplemental Fig. S4: Comparison of drug activity metrics.** **a**, comparison of euclidean distance and classifier performance (AUROC) to describe drug activity. **b**, average AUROC observed and standard deviation of AUROC estimates after 10-fold cross validation. **c**. Hierarchical clustering of active compound effects by pearson correlation. Compound effects were determined by concatenating average morphology profiles from all organoid lines (25 principal components).

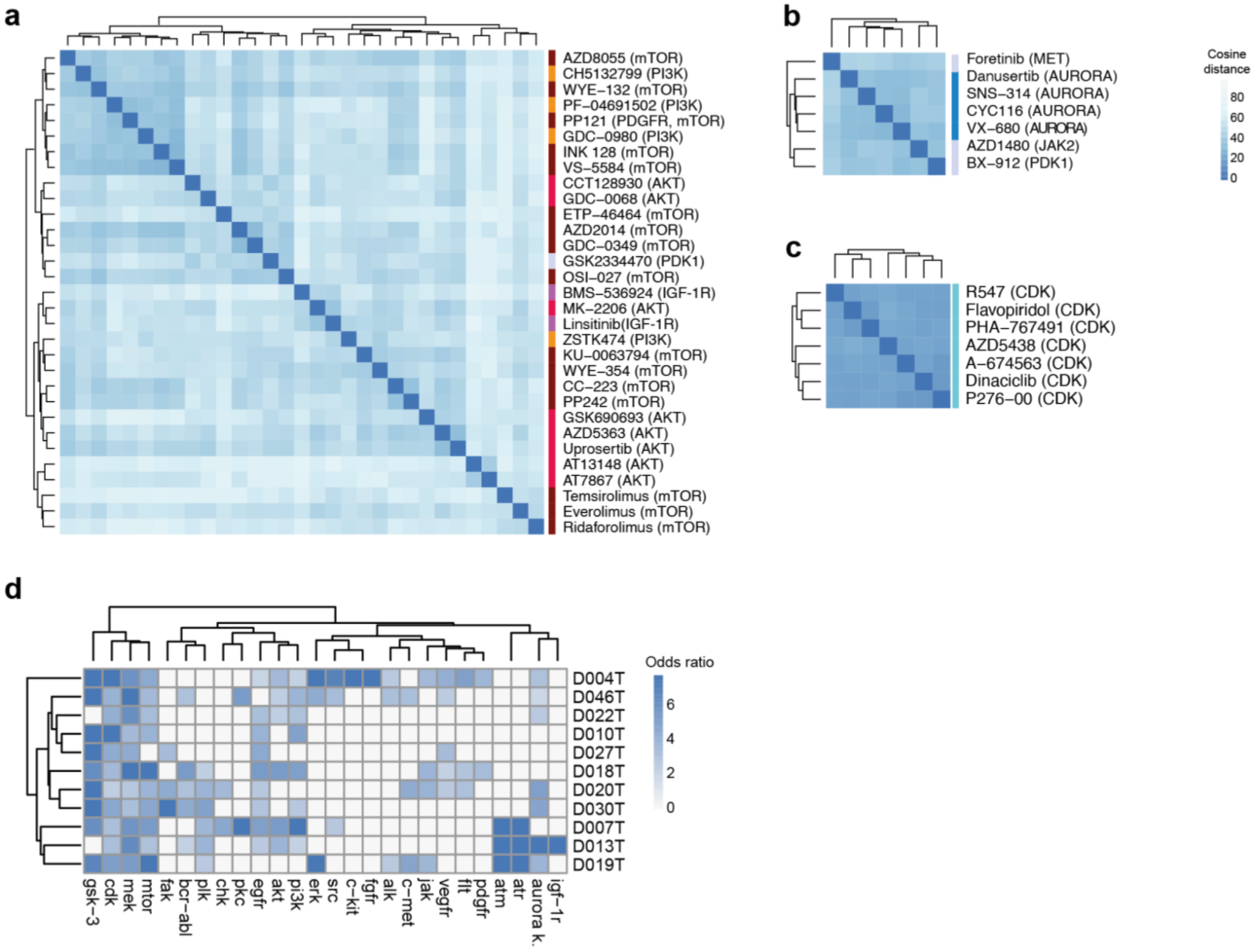


**Supplemental Fig. S5: Selected drug-induced profile clusters** (From Fig. 3d). **a-c**, shown are clusters enriched for PI3K/AKT/mTOR signaling inhibitors, Aurora kinase and CDK inhibitors, respectively. **d**, Significantly enriched targets across profiled PDOs. The cosine distance between drug effect vectors in individual organoid models was clustered, enrichment of annotated molecular targets was tested using Fisher's exact test. Log odds ratios for enriched targets and organoid lines are shown.

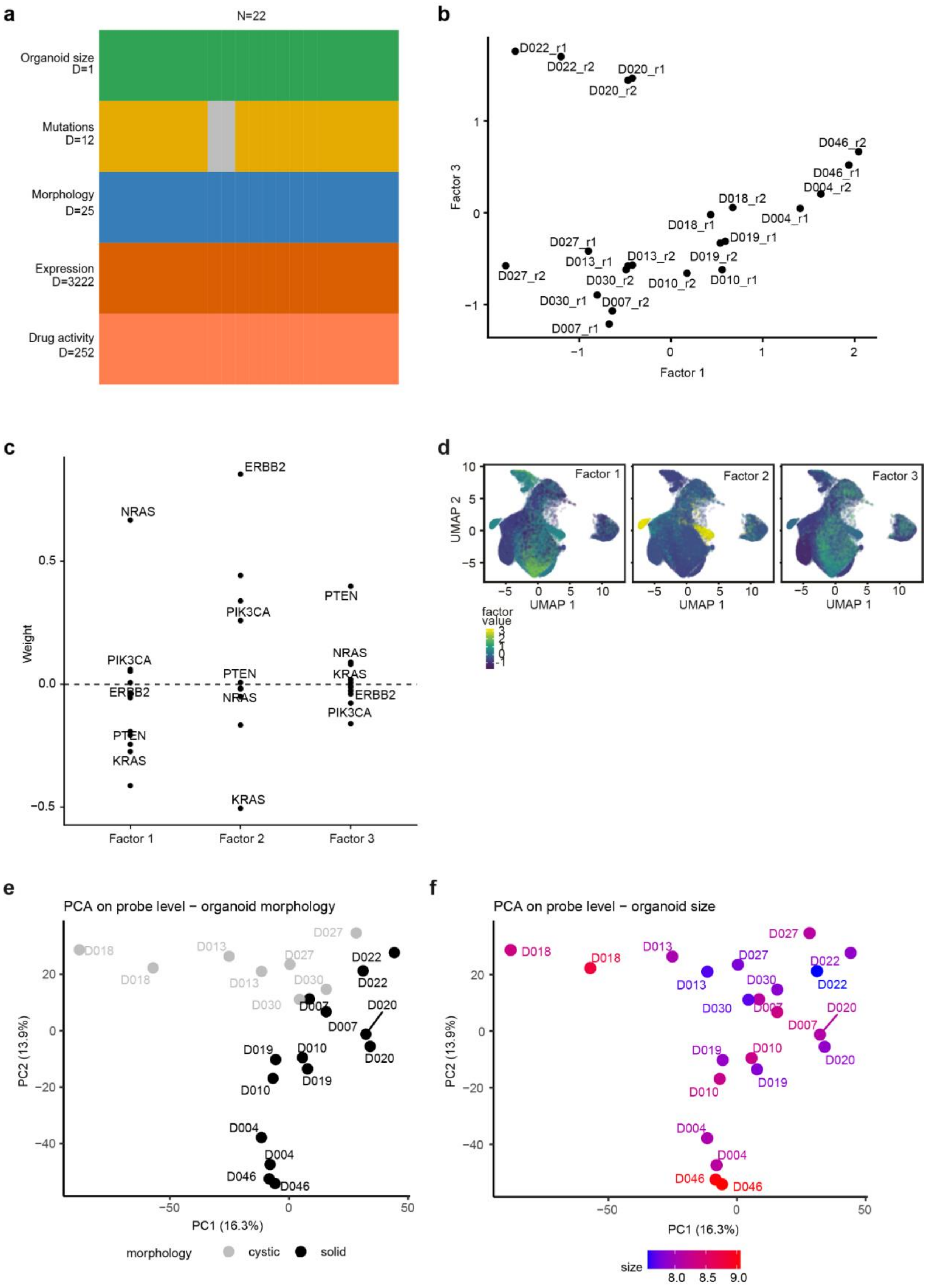
a
N=22
Organoid size
D=1
Mutations
D=12
Morphology
D=25
Expression
D=3222
Drug activity
D=252
b
Factor 3
Factor 1
D022_r1
D022_r2
D020_r1
D020_r2
D046_r2
D046_r1
D004_r2
D004_r1
D018_r2
D018_r1
D019_r1
D019_r2
D027_r1
D027_r2
D013_r1
D013_r2
D030_r2
D030_r1
D010_r2
D010_r1
D007_r2
D007_r1
c
Weight
ERBB2
NRAS
PIK3CA
PTEN
KRAS
Factor 1
Factor 2
Factor 3
d
Factor 1
Factor 2
Factor 3
UMAP 2
UMAP 1
factor value
e
PCA on probe level − organoid morphology
PC2 (13.9%)
PC1 (16.3%)
D018
D013
D027
D030
D022
D007
D020
D019
D010
D004
D046
morphology
cystic
solid
f
PCA on probe level − organoid size
PC2 (13.9%)
PC1 (16.3%)
size
8.0
8.5
9.0

**Supplemental Fig. S6: Multi-omics factor analysis overview. a,** input data matrix showing the five included data modalities and 22 observations (11 lines a' 2 replicates). **b**, Factor 3 scores across unperturbed organoid lines. **c,** Factor loadings for somatic mutations, as identified by amplicon sequencing. **d**, UMAP embedding with organoid lines colored by factor scores. **e-f,** PCA analysis of expression data of unperturbed organoid lines. Two biological replicates were performed of all organoid lines. Colors represent organoid morphology (cystic vs. solid, visual inspection, e) and organoid size (f).

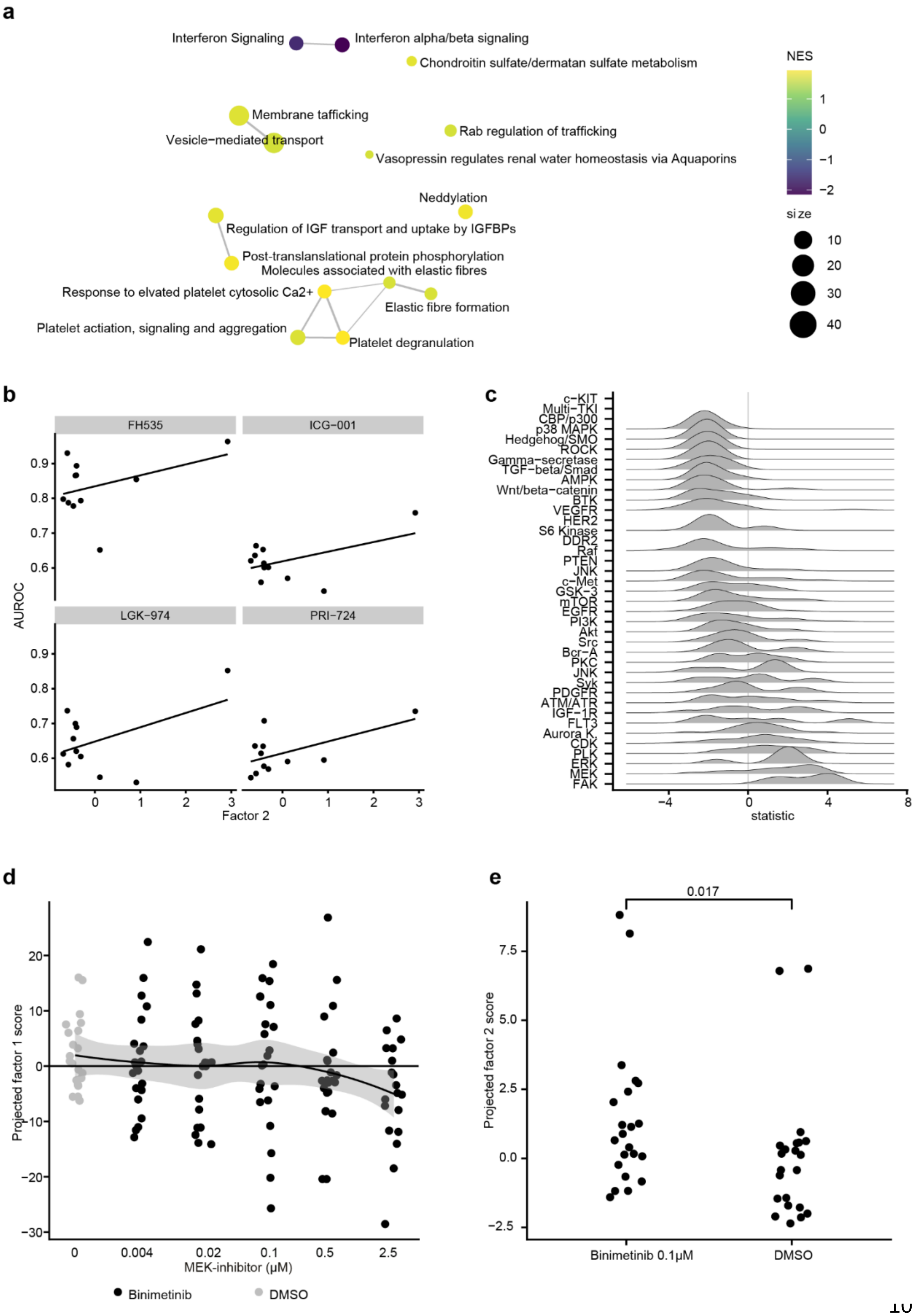
a
Interferon Signaling
Interferon alpha/beta signaling
Chondroitin sulfate/dermatan sulfate metabolism
Membrane tafficking
Vesicle−mediated transport
Rab regulation of trafficking
Vasopressin regulates renal water homeostasis via Aquaporins
Neddylation
Regulation of IGF transport and uptake by IGFBPs
Post-translanslational protein phosphorylation
Molecules associated with elastic fibres
Response to elvated platelet cytosolic Ca2+
Elastic fibre formation
Platelet actiation, signaling and aggregation
Platelet degranulation
NES
size
b
FH535
ICG−001
LGK−974
PRI−724
AUROC
Factor 2
c
c−KIT
Multi−TKI
CBP/p300
p38 MAPK
Hedgehog/SMO
ROCK
Gamma−secretase
TGF−beta/Smad
AMPK
Wnt/beta−catenin
BTK
VEGFR
HER2
S6 Kinase
DDR2
Raf
PTEN
JNK
c−Met
GSK−3
mTOR
EGFR
PI3K
Akt
Src
Bcr−A
PKC
JNK
Syk
PDGFR
ATM/ATR
IGF−1R
FLT3
Aurora K.
CDK
PLK
ERK
MEK
FAK
statistic
d
Projected factor 1 score
MEK-inhibitor (μM)
Binimetinib
DMSO
e
0.017
Projected factor 2 score
Binimetinib 0.1μM
DMSO

**Supplemental Fig. S7: Factor 2 defined by organoid LGR5 stem cell identity. a**, Reactome pathway enrichment results for factor 2 gene expression loadings. Colorscale corresponds to normalized enrichment score (NES), where a positive value stands for an enrichment and a negative value for a depletion. Size of the nodes represent gene set size, edges correspond to gene set overlap **b,** Relationship of Wnt pathway inhibitor activity with factor 2 score. **c,** Approximated factor 2 shifts measured in organoids treated with small molecule inhibitors when accounting for line specific differences. Shifts are expressed as t statistics for the model (factor 2 ~ drug + line). **d,** Dose-dependent changes in factor 1 (IGFR1 signaling) scores after treatment with the MEK inhibitor binimitenib across organoid lines. **e,** Factor 2 scores for organoids treated with 1uM binimetinib and DMSO control, two-sided Wilcoxon rank sum test.

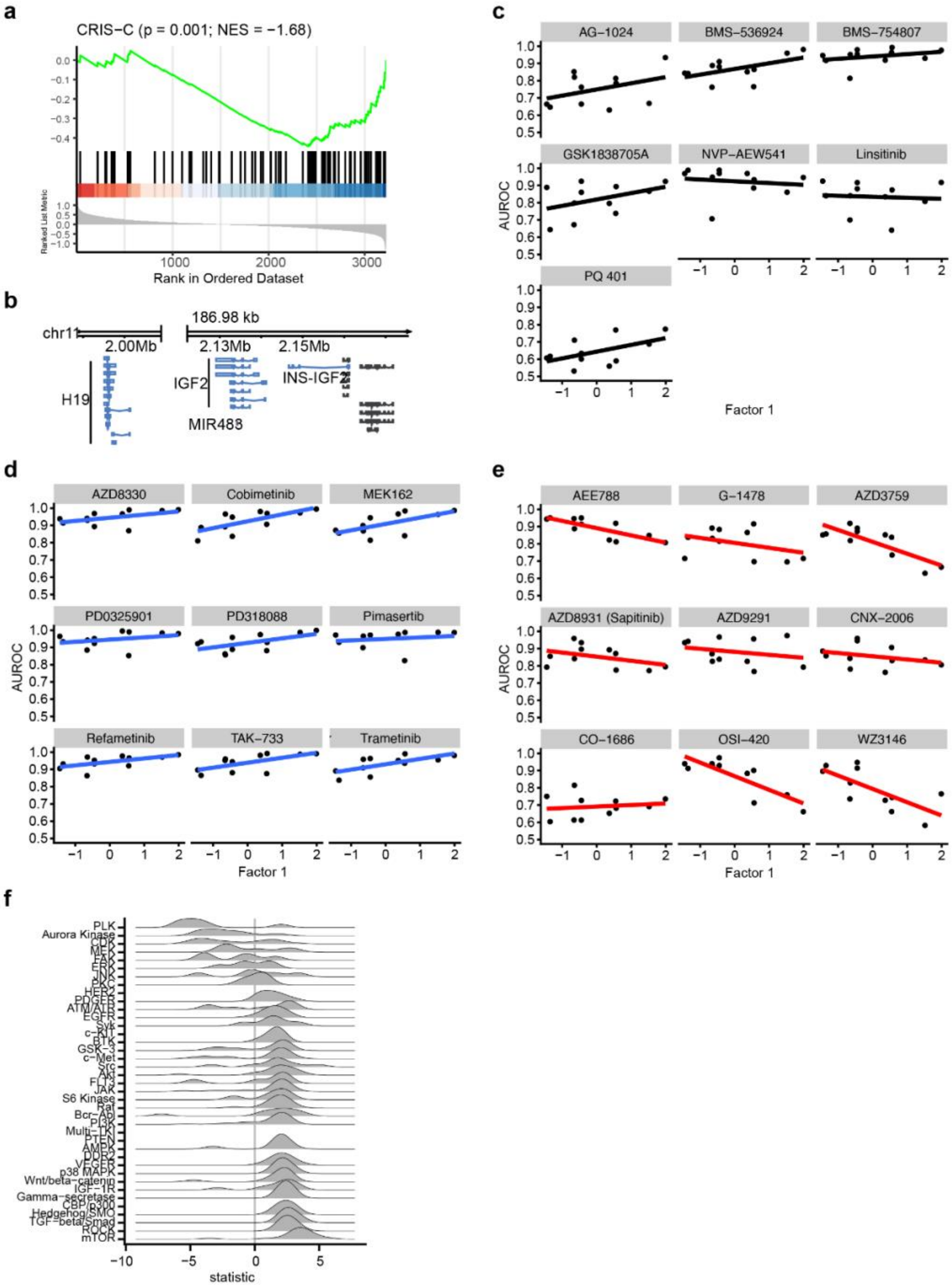


**Supplemental Fig. S8: Factor 1 defined by IGF1R receptor signaling.** **a**, gene set enrichment analysis for CRIS signature C on factor 1 gene expression loadings. **b,** Genomic track plot for IGF2, H19 and INS-IGF2. **c-e,** Relationship of IGF1R-, MEK- and EGFR-inhibitor activity with factor 1 scores, respectively. Shown are 9 randomly sampled small molecule inhibitors. **f,** Approximated factor 1 shifts measured in organoids treated with small molecule inhibitors when accounting for line specific differences. Shifts are expressed as t statistics for the model (factor 1 ~ drug + line).
