## Supplemental Tables for "The drug-induced phenotypic landscape of colorectal cancer organoids"

### Table S1: Patient and organoid characteristics

| PDO line | PDO medium | PDO growth | Patient age | Patient sex | Tumor location | Tumor stage (UICC) | Tumor grade (WHO) | Tumor MSI status | Tumor treatment |
|---|---|---|---|---|---|---|---|---|---|
| **D004T** | ENA | good | 62 | f | rectum | 3 | 2 | MSS | neoadj RCTX, OP, adj. CTX |
| **D007T** | ENA | good | 33 | m | rectum | 3 | 2 | MSS | neoadj RCTX, OP, adj. CTX |
| **D010T** | ENA | good | 58 | m | sigm | 3 | 2 | MSS | OP |
| **D013T** | ENAS | good | 84 | f | rectum | 1 | 2 | n.d. | endoscopic resection |
| **D015T** | ENA | good | 88 | m | desc | 2 | 3 | MSS | OP |
| **D018T** | ENA | good | 84 | m | sigm | 2 | 2 | MSS | OP |
| **D019T** | ENA | good | 50 | f | sigm | 1 | 2 | MSS | OP |
| **D020T** | ENA | good | 86 | m | rectum | 1 | 3 | MSS | OP |
| **D021T** | ENA | medium | 68 | m | rectum | 1 | n.d. | n.d. | OP |
| **D022T** | ENA | good | 64 | m | asc | 4 | 2 | MSS | CTX |
| **D027T** | ENA | good | 65 | m | rectum | 4 | 2 | MSS | CTX |
| **D030T** | ENA | good | 71 | f | asc | 2 | 2 | MSS | OP, adj CTX |
| **D046T** | ENA | good | 51 | f | rectum | 3 | 2 | MSS | neoadj RCTX, OP, adj CTX |

Abbreviations: f, female; m, male; sigm, sigmoid; desc, descending colon; asc, ascending colon; MSI, microsatelite instability, MSS, microsatellite stable; OP, operation; neoadj; neoadjuvant; RCTX, radiochemotherapy; CTX, chemotherapy; ENA, basal medium + EGF, Noggin and A83-01 (ALK-inhibitor); ENAS, ENA + SB202190 (p38 inhibitor).

### Table S2: Mutations in organoids detected by amplicon sequencing

| SAMPLE | SYMBOL | Protein_position | Amino_acids | Consequence |
|---|---|---|---|---|
| P004T | NRAS | 61 | Q/L | missense_variant |
| P004T | PIK3CA | 546 | Q/P | missense_variant |
| P007T | AKT1 | 17 | E/K | missense_variant |
| P007T | APC | 564 | R/* | stop_gained |
| P007T | KRAS | 12 | G/D | missense_variant |
| P010T | KRAS | 13 | G/D | missense_variant |
| P010T | TP53 | - | - | splice_acceptor_variant |
| P013T | APC | 876 | R/* | stop_gained |
| P013T | APC | 1450 | R/* | stop_gained |
| P013T | ERBB2 | 310 | S/F | missense_variant |
| P013T | ERBB2 | 769 | D/Y | missense_variant |
| P013T | PIK3CA | 1004 | M/I | missense_variant |
| P013T | TP53 | 273 | R/C | missense_variant |
| P015T | ATM | 2809 | Q/X | frameshift_variant |
| P015T | B2M | 14-15 | SL/X | frameshift_variant |
| P015T | CTNNB1 | 34 | G/V | missense_variant |
| P015T | FBXW7 | 465 | R/C | missense_variant |
| P015T | KRAS | 13 | G/D | missense_variant |
| P015T | PTEN | 160 | T/I | missense_variant |
| P015T | TCF7L2 | 465 | R/X | frameshift_variant |
| P019T | APC | 1378 | Q/* | stop_gained |
| P019T | ARID1A | 1844 | W/* | stop_gained |
| P019T | KRAS | 146 | A/T | missense_variant |
| P019T | RNF43 | 225 | R/C | missense_variant |
| P019T | SMAD4 | 361 | R/C | missense_variant |
| P019T | TP53 | 234 | Y/N | missense_variant |
| P020T | AMER1 | 511 | P/A | missense_variant |
| P020T | KRAS | 12 | G/V | missense_variant |
| P020T | SMAD4 | 102 | P/R | missense_variant |
| P021T | APC | 1356 | S/* | stop_gained |
| P021T | GNAS | 844 | R/H | missense_variant |
| P021T | KRAS | 12 | G/D | missense_variant |
| P022T | APC | 232 | R/* | stop_gained |
| P022T | KRAS | 12 | G/C | missense_variant |
| P022T | PTEN | 137 | A/V | missense_variant |
| P022T | TP53 | 132 | K/R | missense_variant |
| P027T | AMER1 | 373 | A/G | missense_variant |
| P027T | APC | 216 | R/* | stop_gained |
| P027T | TP53 | 282 | R/W | missense_variant |
| P030T | KRAS | 146 | A/T | missense_variant |
| P030T | PIK3CA | 420 | C/R | missense_variant |
| P030T | SMAD4 | 537 | D/A | missense_variant |
| P046T | NRAS | 61 | Q/L | missense_variant |

### Table S3: Clinical cancer compound library

| Compound Name | Target | Category | Phase | Max conc (µM) | Min conc (µM) |
|---|---|---|---|---|---|
| 17AAG | HSP90 | Targeted | Phase III | 12,5 | 0,02 |
| Afatinib | ErbB-Receptors | Targeted | Clinical use | 100 | 0,16 |
| Alisertib | Aurora A | Targeted | Phase II/III | 25 | 0,04 |
| Alpelisib | PIK3a | Targeted | Phase I | 25 | 0,04 |
| Axitinib | PDGFR, KIT, VEGFR | Targeted | Clinical use | 75 | 0,12 |
| AZD 4547 | FGFR | Targeted | Phase II/III | 25 | 0,04 |
| AZD 5363 | AKT | Targeted | Phase II | 25 | 0,04 |
| Bexaroten | Retinoid-Receptor | Targeted | Clinical use | 20 | 0,032 |
| Binimetinib | NRAS, MEK | Targeted | Phase III | 25 | 0,04 |
| Birinapant | SMAC | Targeted | Phase II | 25 | 0,04 |
| Bortezomib | Proteasom | Targeted | Clinical use | 2,5 | 0,004 |
| Cabozantinib | VEGF, MET, RET | Targeted | Clinical use | 100 | 0,16 |
| Crizotinib | ALK | Targeted | Clinical use | 12,5 | 0,02 |
| Dabrafenib | BRAF | Targeted | Clinical use | 50 | 0,08 |
| Dasatinib | DDR2, BCR-ABL, SRC, KIT, PDGFR | Targeted | Clinical use | 100 | 0,16 |
| Defactinib | FAK | Targeted | Phase II | 7,5 | 0,012 |
| Erlotinib | EGFR | Targeted | Clinical use | 100 | 0,16 |
| Everolimus | mTOR | Targeted | Clinical use | 5 | 0,008 |
| Gefitinib | EGFR | Targeted | Clinical use | 100 | 0,16 |
| GSK2636771 | PTEN | Targeted | Phase I/II | 25 | 0,04 |
| Idelalisib | PIK3δ | Targeted | Clinical use | 100 | 0,16 |
| Lapatinib | HER2 and EGFR | Targeted | Clinical use | 100 | 0,16 |
| LGK974 | Porcupine | Targeted | Phase I | 100 | 0,16 |
| MK-1775 | Wee1 | Targeted | Phase II | 25 | 0,04 |
| MK-8776 | Chk1 | Targeted | Phase II | 7,5 | 0,012 |
| Napabucasin | STAT3 | Targeted | Phase III | 25 | 0,04 |
| Nutlin3a | p53 | Targeted | Preclinical | 100 | 0,16 |
| Olaparib | PARP | Targeted | Clinical use | 100 | 0,16 |
| Palbociclib | CDK | Targeted | Clinical use | 5 | 0,008 |
| Panobinostat | HDAC | Targeted | Clinical use | 2,5 | 0,004 |
| Pazopanib | c-KIT, FGFR, PDGFR and VEGFR | Targeted | Clinical use | 100 | 0,16 |
| Ponatinib | BEGFR, PDGFR, FGFR, EPH recep | Targeted | Clinical use | 25 | 0,04 |
| PRI-724 | CBP/p300 | Targeted | Phase I-II | 75 | 0,12 |
| Regorafenib | VEGFR-1/2/3, TIE-2, KIT, RET, RAF | Targeted | Clinical use | 100 | 0,16 |
| Ruxolitinib | JAK1 and JAK2 | Targeted | Clinical use | 100 | 0,16 |
| Sonidegib | SMO | Targeted | Clinical use | 100 | 0,16 |
| Sorafenib | PDGFR, KIT, VEGFR | Targeted | Clinical use | 100 | 0,16 |
| Sunitinib | Multi-TKI | Targeted | Clinical use | 50 | 0,08 |
| Taselisib | PI3K | Targeted | Phase III | 25 | 0,04 |
| Trametinib | Mek1/2 | Targeted | Clinical use | 2,5 | 0,004 |
| Venetoclax | Bcl-2 | Targeted | Clinical use | 25 | 0,04 |
| Vismodegib | SMO | Targeted | Clinical use | 100 | 0,16 |
| Volasertib | PLK1 | Targeted | Phase II | 25 | 0,04 |
| Vorinostat | HDAC | Targeted | Clinical use | 100 | 0,16 |
| VX-702 | p38 MAPK | Targeted | Phase II | 25 | 0,04 |
| YM155 | Survivin | Tagreted | Phase II | 5 | 0,008 |
| 5-FU | Antimetabolite | Chemotherapy | Clinical use | 100 | 0,16 |
| Bleomycin | DNA-Damage | Chemotherapy | Clinical use | 50 | 0,08 |
| Dacarbazine | Alkylating | Chemotherapy | Clinical use | 12,5 | 0,02 |
| Docetaxel | Mikrotubuli | Chemotherapy | Clinical use | 2,5 | 0,004 |
| Doxorubicin | Intercalating, mRNA synthesis | Chemotherapy | Clinical use | 50 | 0,08 |
| Etoposid | Topoisomerase | Chemotherapy | Clinical use | 100 | 0,16 |
| Fludarabine | Antimetabolite | Chemotherapy | Clinical use | 12,5 | 0,02 |
| Gemcitabine | DMA-Synthesis | Chemotherapy | Clinical use | 12,5 | 0,02 |
| Irinotecan / SN-38 | Topoisomerase | Chemotherapy | Clinical use | 25 | 0,04 |
| Methotrexate | Antimetabolite | Chemotherapy | Clinical use | 50 | 0,08 |
| Mitomycin C | Alkylating | Chemotherapy | Clinical use | 12,5 | 0,02 |
| Oxaliplatin | Alkylating | Chemotherapy | Clinical use | 75 | 0,12 |
| Paclitaxel | Mikrotubuli | Chemotherapy | Clinical use | 2,5 | 0,004 |
| Trifluoridin/Tipiracil | Antimetabolite | Chemotherapy | Clinical use | 55 | 0,088 |
| Vinblastin | Mikrotubuli | Chemotherapy | Clinical use | 2,5 | 0,004 |
| Miglitol | Antidiabetic | Metabolism | Clinical use | 12,5 | 0,02 |
| Phenformin | Antidiabetic | Metabolism | Clinical use | 100 | 0,16 |

### Table S4: Kinase-Stemcell (Ki-Stem) compound library

| Compound name | Target | Pathway | conc (uM) |
|---|---|---|---|
| 1-Azakenpaullone | GSK-3 | PI3K/Akt/mTOR | 7,5 |
| 10058-F4 | c-Myc | Cell Cycle | 7,5 |
| 3-Methyladenine | PI3K | PI3K/Akt/mTOR | 7,5 |
| 7,8-Dihydroxyflavone | Trk receptor | Protein Tyrosine Kinase | 7,5 |
| A-674563 | Akt, CDK, PKA | PI3K/Akt/mTOR | 7,5 |
| A-769662 | AMPK | PI3K/Akt/mTOR | 7,5 |
| A66 | PI3K | PI3K/Akt/mTOR | 7,5 |
| AC480 (BMS-599626) | HER2 | Neuronal Signaling | 7,5 |
| Acadesine | AMPK | PI3K/Akt/mTOR | 7,5 |
| AEE788 (NVP-AEE788) | EGFR, Flt, VEGFR, HER2 | Protein Tyrosine Kinase | 7,5 |
| Afatinib (BIBW2992) | EGFR | Protein Tyrosine Kinase | 7,5 |
| AG-1024 | IGF-1R | Protein Tyrosine Kinase | 7,5 |
| AG-1478 (Tyrphostin AG-1478) | EGFR | Protein Tyrosine Kinase | 7,5 |
| AG-18 | EGFR | Protein Tyrosine Kinase | 7,5 |
| AG-490 (Tyrphostin B42) | JAK, EGFR | Others | 7,5 |
| Akti-1/2 | Akt | PI3K/Akt/mTOR | 7,5 |
| Alisertib (MLN8237) | Aurora Kinase | Others | 7,5 |
| AMG 337 | c-Met | Protein Tyrosine Kinase | 7,5 |
| AMG-458 | c-Met | Protein Tyrosine Kinase | 7,5 |
| AMG-900 | Aurora Kinase | Cell Cycle | 7,5 |
| AMG319 | PI3K | PI3K/Akt/mTOR | 7,5 |
| Amuvatinib (MP-470) | c-Met, c-Kit, PDGFR, Flt, c-RET | Protein Tyrosine Kinase | 7,5 |
| ANA-12 | Trk receptor | Protein Tyrosine Kinase | 7,5 |
| Anacardic Acid | Histone Acetyltransferase | Epigenetics | 7,5 |
| AP26113 | ALK | Protein Tyrosine Kinase | 7,5 |
| Apatinib | VEGFR | Protein Tyrosine Kinase | 7,5 |
| AR-A014418 | GSK-3 | PI3K/Akt/mTOR | 7,5 |
| AR-A014418 | GSK-3 | PI3K/Akt/mTOR | 7,5 |
| AS-252424 | PI3K | PI3K/Akt/mTOR | 7,5 |
| AS-604850 | PI3K | PI3K/Akt/mTOR | 7,5 |
| Asiatic Acid | p38 MAPK | MAPK | 7,5 |
| ASP3026 | ALK | Protein Tyrosine Kinase | 7,5 |
| AST-1306 | EGFR | Protein Tyrosine Kinase | 7,5 |
| Astragaloside A | others | TGF-beta/Smad | 7,5 |
| AT13148 | Akt | PI3K/Akt/mTOR | 7,5 |
| AT7519 | CDK | Cell Cycle | 7,5 |
| AT7867 | Akt, S6 kinase | PI3K/Akt/mTOR | 7,5 |
| AT9283 | Bcr-Abl, JAK, Aurora Kinase | Others | 7,5 |
| Aurora A Inhibitor I | Aurora Kinase | Cell Cycle | 7,5 |
| Avagacestat (BMS-708163) | Gamma-secretase | Proteases | 7,5 |
| AVL-292 | BTK | Angiogenesis | 7,5 |
| Axitinib | VEGFR, PDGFR, c-Kit | Protein Tyrosine Kinase | 7,5 |
| AZ 628 | Raf | MAPK | 7,5 |
| AZ 960 | JAK | JAK/STAT | 7,5 |
| AZ20 | ATM/ATR | PI3K/Akt/mTOR | 7,5 |
| AZD1080 | GSK-3 | PI3K/Akt/mTOR | 7,5 |
| AZD1208 | Pim | JAK/STAT | 7,5 |
| AZD1480 | JAK | JAK/STAT | 7,5 |
| AZD1480 | JAK | JAK/STAT | 7,5 |
| AZD2014 | mTOR | PI3K/Akt/mTOR | 7,5 |
| AZD2858 | GSK-3 | PI3K/Akt/mTOR | 7,5 |
| AZD2932 | PDGFR | Protein Tyrosine Kinase | 7,5 |
| AZD3463 | ALK | Protein Tyrosine Kinase | 7,5 |
| AZD3759 | EGFR | Protein Tyrosine Kinase | 7,5 |
| AZD4547 | FGFR | Angiogenesis | 7,5 |
| AZD5363 | Akt | PI3K/Akt/mTOR | 7,5 |
| AZD5438 | CDK | Cell Cycle | 7,5 |
| AZD6482 | PI3K | PI3K/Akt/mTOR | 7,5 |
| AZD6738 | ATM/ATR | PI3K/Akt/mTOR | 7,5 |
| AZD7762 | Chk | Cell Cycle | 7,5 |
| AZD8055 | mTOR | PI3K/Akt/mTOR | 7,5 |
| AZD8330 | MEK | MAPK | 7,5 |
| AZD8931 (Sapitinib) | EGFR, HER2 | Protein Tyrosine Kinase | 7,5 |
| AZD9291 | EGFR | Protein Tyrosine Kinase | 7,5 |
| Barasertib (AZD1152-HQPA) | Aurora Kinase | Others | 7,5 |
| Baricitinib (LY3009104, INCB028 | JAK | Epigenetics | 7,5 |
| BAY 11-7082 | I_B/IKK | NF-_B | 7,5 |
| Bay 11-7085 | I_B/IKK | NF-_B | 7,5 |
| BGT226 (NVP-BGT226) | PI3K, mTOR | PI3K/Akt/mTOR | 7,5 |
| BI 2536 | PLK | Others | 7,5 |
| BI-78D3 | JNK | MAPK | 7,5 |
| BI-D1870 | S6 Kinase | PI3K/Akt/mTOR | 7,5 |
| Bikinin | GSK-3 | PI3K/Akt/mTOR | 7,5 |
| BIO | GSK-3 | PI3K/Akt/mTOR | 7,5 |
| BIRB 796 (Doramapimod) | p38 MAPK | MAPK | 7,5 |

| | | | |
|---|---|---|---|
| BIX 02188 | MEK | MAPK | 7,5 |
| BKM120 (NVP-BKM120, Buparli | PI3K | PI3K/Akt/mTOR | 7,5 |
| BLZ945 | CSF-1R | Protein Tyrosine Kinase | 7,5 |
| BMS-265246 | CDK | Cell Cycle | 7,5 |
| BMS-345541 | I_B/IKK | NF-_B | 7,5 |
| BMS-536924 | IGF-1R | Protein Tyrosine Kinase | 7,5 |
| BMS-582949 | p38 MAPK | MAPK | 7,5 |
| BMS-754807 | IGF-1R | Others | 7,5 |
| BMS-777607 | c-Met | Protein Tyrosine Kinase | 7,5 |
| BMS-794833 | c-Met, VEGFR | Protein Tyrosine Kinase | 7,5 |
| BMS-833923 | Hedgehog/Smoothened | GPCR & G Protein | 7,5 |
| Bosutinib (SKI-606) | Src | Angiogenesis | 7,5 |
| Brivanib (BMS-540215) | VEGFR, FGFR | GPCR & G Protein | 7,5 |
| Brivanib Alaninate (BMS-582664 | VEGFR, FGFR | Others | 7,5 |
| BS-181 HCl | CDK | Cell Cycle | 7,5 |
| Butein | EGFR | Protein Tyrosine Kinase | 7,5 |
| BX-795 | PDK-1, IKK | PI3K/Akt/mTOR | 7,5 |
| BX-912 | PDK-1 | PI3K/Akt/mTOR | 7,5 |
| BYL719 | PI3K | PI3K/Akt/mTOR | 7,5 |
| Cabozantinib (XL184, BMS-907 | VEGFR, c-Met, Flt, Tie-2, c-Kit | Others | 7,5 |
| CAL-101 (Idelalisib, GS-1101) | PI3K | PI3K/Akt/mTOR | 7,5 |
| CAY10505 | PI3K | PI3K/Akt/mTOR | 7,5 |
| CC-223 | mTOR | PI3K/Akt/mTOR | 7,5 |
| CCT128930 | Akt | PI3K/Akt/mTOR | 7,5 |
| Cediranib (AZD2171) | VEGFR, Flt | Protein Tyrosine Kinase | 7,5 |
| CEP-32496 | Raf | MAPK | 7,5 |
| CEP-33779 | JAK | JAK/STAT | 7,5 |
| CGI1746 | BTK | Angiogenesis | 7,5 |
| CGK 733 | ATM/ATR | DNA Damage | 7,5 |
| CH5132799 | PI3K, mTOR | PI3K/Akt/mTOR | 7,5 |
| CHIR-124 | Chk | Cell Cycle | 7,5 |
| CHIR-98014 | GSK-3 | PI3K/Akt/mTOR | 7,5 |
| CHIR-99021 (CT99021) | GSK-3 | PI3K/Akt/mTOR | 7,5 |
| Chrysophanic Acid | EGFR, mTOR | Protein Tyrosine Kinase | 7,5 |
| CNX-2006 | EGFR | Protein Tyrosine Kinase | 7,5 |
| CNX-774 | BTK | Angiogenesis | 7,5 |
| CO-1686 (AVL-301) | EGFR | Protein Tyrosine Kinase | 7,5 |
| Cobimetinib (GDC-0973, RG742 | MEK | MAPK | 7,5 |
| CP-673451 | PDGFR | Protein Tyrosine Kinase | 7,5 |
| CP-724714 | EGFR, HER2 | Protein Tyrosine Kinase | 7,5 |
| Crenolanib (CP-868596) | PDGFR | Protein Tyrosine Kinase | 7,5 |
| Crizotinib (PF-02341066) | c-Met, ALK | Others | 7,5 |
| CUDC-101 | HDAC, EGFR, HER2 | Epigenetics | 7,5 |
| CUDC-907 | HDAC, PI3K | Cytoskeletal Signaling | 7,5 |
| CX-6258 HCl | Pim | JAK/STAT | 7,5 |
| CYC116 | Aurora Kinase, VEGFR | Cell Cycle | 7,5 |
| CYT387 | JAK | JAK/STAT | 7,5 |
| CZC24832 | PI3K_ | PI3K/Akt/mTOR | 7,5 |
| D 4476 | CK | Metabolism | 7,5 |
| Dabrafenib (GSK2118436) | Raf | MAPK | 7,5 |
| Dacomitinib (PF299804, PF299) | EGFR | Protein Tyrosine Kinase | 7,5 |
| Danusertib (PHA-739358) | Aurora Kinase, FGFR, Bcr-Abl, | Others | 7,5 |
| Daphnetin | PKA/PKC/EGFR | DNA Damage | 7,5 |
| DAPT (GSI-IX) | Gamma-secretase, Beta Amyloi | Proteases | 7,5 |
| DASA-58 | PKM2 | Others | 7,5 |
| Dasatinib | Src, Bcr-Abl, c-Kit | Angiogenesis | 7,5 |
| DCC-2036 (Rebastinib) | Bcr-Abl | Angiogenesis | 7,5 |
| DDR1-IN-1 | DDR(receptor tyrosine kinase) | Others | 7,5 |
| Decernotinib (VX-509) | JAK | JAK/STAT | 7,5 |
| Degrasyn (WP1130) | DUB, Bcr-Abl | Angiogenesis | 7,5 |
| Dinaciclib (SCH727965) | CDK | Cell Cycle | 7,5 |
| Dovitinib (TKI-258, CHIR-258) | c-Kit, FGFR, Flt, VEGFR, PDGI | Angiogenesis | 7,5 |
| Dovitinib Dilactic Acid | Flt, FGFR, PDGFR, VEGFR, c- | Angiogenesis | 7,5 |
| EHop-016 | Rac | Cell Cycle | 7,5 |
| ENMD-2076 | Flt, Aurora Kinase, VEGFR | Angiogenesis | 7,5 |
| Entrectinib (RXDX-101) | Trk receptor | Protein Tyrosine Kinase | 7,5 |
| Enzastaurin (LY317615) | PKC | Neuronal Signaling | 7,5 |
| ERK5-IN-1 | ERK | MAPK | 7,5 |
| ETC-1002 | AMPK | PI3K/Akt/mTOR | 7,5 |
| ETP-46464 | mTOR | PI3K/Akt/mTOR | 7,5 |
| Everolimus (RAD001) | mTOR | Others | 7,5 |
| Fasudil (HA-1077) HCl | ROCK | Cell Cycle | 7,5 |
| FH535 | Wnt/beta-catenin | Stem Cells & Wnt | 7,5 |
| FIIN-2 | FGFR | Protein Tyrosine Kinase | 7,5 |
| Filgotinib (GLPG0634) | JAK | JAK/STAT | 7,5 |
| Fingolimod (FTY720) HCl | S1P Receptor, Bcr-Abl, PKC | GPCR & G Protein | 7,5 |
| Flavopiridol (Alvocidib) | CDK | Cell Cycle | 7,5 |
| Foretinib (GSK1363089) | c-Met, VEGFR | Others | 7,5 |

| | | | |
|---|---|---|---|
| Fostamatinib (R788) | Syk | Angiogenesis | 7,5 |
| FRAX597 | PAK | Cytoskeletal Signaling | 7,5 |
| G-749 | FLT3 | Angiogenesis | 7,5 |
| GDC-0068 | Akt | PI3K/Akt/mTOR | 7,5 |
| GDC-0349 | mTOR | PI3K/Akt/mTOR | 7,5 |
| GDC-0879 | Raf | Others | 7,5 |
| GDC-0941 | PI3K | Metabolism | 7,5 |
| GDC-0980 (RG7422) | mTOR, PI3K | PI3K/Akt/mTOR | 7,5 |
| Gefitinib (ZD1839) | EGFR | Protein Tyrosine Kinase | 7,5 |
| Genistein | EGFR | Protein Tyrosine Kinase | 7,5 |
| GF109203X | PKC | TGF-beta/Smad | 7,5 |
| GF109203X | PKC | TGF-beta/Smad | 7,5 |
| GNE-0877 | leucine-rich repeat kinase 2 (LR | Autophagy | 7,5 |
| GNE-7915 | LRRK | Autophagy | 7,5 |
| GNE-9605 | LRRK2 | Autophagy | 7,5 |
| GNF-2 | Bcr-Abl | Angiogenesis | 7,5 |
| GNF-5 | Bcr-Abl | Angiogenesis | 7,5 |
| Go 6983 | PKC | TGF-beta/Smad | 7,5 |
| Golvatinib (E7050) | c-Met, VEGFR | Protein Tyrosine Kinase | 7,5 |
| GSK1838705A | IGF-1, ALK | Protein Tyrosine Kinase | 7,5 |
| GSK1904529A | IGF-1R | Others | 7,5 |
| GSK2126458 (GSK458) | PI3K, mTOR | PI3K/Akt/mTOR | 7,5 |
| GSK2292767 | PI3K | PI3K/Akt/mTOR | 7,5 |
| GSK2334470 | PDK-1 | PI3K/Akt/mTOR | 7,5 |
| GSK2578215A | LRRK2 | Autophagy | 7,5 |
| GSK2636771 | PI3K | PI3K/Akt/mTOR | 7,5 |
| GSK429286A | ROCK | Cell Cycle | 7,5 |
| GSK461364 | PLK | Cell Cycle | 7,5 |
| GSK621 | AMPK | PI3K/Akt/mTOR | 7,5 |
| GSK650394 | SGK | Others | 7,5 |
| GSK690693 | Akt | Others | 7,5 |
| GW441756 | Trk Receptor | Protein Tyrosine Kinase | 7,5 |
| GW5074 | Raf | MAPK | 7,5 |
| GW788388 | TGF-beta/Smad | TGF-beta/Smad | 7,5 |
| GZD824 | Bcr-Abl | Angiogenesis | 7,5 |
| H 89 2HCl | S6 Kinase | PI3K/Akt/mTOR | 7,5 |
| Hesperadin | Aurora Kinase | Cell Cycle | 7,5 |
| Hesperetin | Histamine Receptor | TGF-beta/Smad | 7,5 |
| HMN-214 | PLK | Cell Cycle | 7,5 |
| HO-3867 | STAT | JAK/STAT | 7,5 |
| Honokiol | Akt, MEK | PI3K/Akt/mTOR | 7,5 |
| HS-173 | PI3K | PI3K/Akt/mTOR | 7,5 |
| HTH-01-015 | AMPK | PI3K/Akt/mTOR | 7,5 |
| Ibrutinib (PCI-32765) | Src | Angiogenesis | 7,5 |
| ICG-001 | Wnt/beta-catenin | Stem Cells & Wnt | 7,5 |
| Icotinib | EGFR | Protein Tyrosine Kinase | 7,5 |
| IKK-16 (IKK Inhibitor VII) | IKK | NF-_B | 7,5 |
| IM-12 | GSK-3 | PI3K/Akt/mTOR | 7,5 |
| Imatinib Mesylate (STI571) | PDGFR, c-Kit, Bcr-Abl | Protein Tyrosine Kinase | 7,5 |
| IMD 0354 | IKK | NF-_B | 7,5 |
| Indirubin | GSK-3 | PI3K/Akt/mTOR | 7,5 |
| INK 128 (MLN0128) | mTOR | PI3K/Akt/mTOR | 7,5 |
| IPA-3 | PAK | Cytoskeletal Signaling | 7,5 |
| IPI-145 (INK1197) | PI3K | Angiogenesis | 7,5 |
| IWP-L6 | Wnt/beta-catenin | Stem Cells & Wnt | 7,5 |
| IWR-1-endo | Wnt/beta-catenin | Stem Cells & Wnt | 7,5 |
| JNJ-38877605 | c-Met | Others | 7,5 |
| JNJ-7706621 | CDK, Aurora Kinase | Cell Cycle | 7,5 |
| JNK Inhibitor IX | JNK | MAPK | 7,5 |
| JNK-IN-8 | JNK | MAPK | 7,5 |
| K02288 | TGF-beta/Smad | TGF-beta/Smad | 7,5 |
| Ki8751 | VEGFR, c-Kit, PDGFR | Protein Tyrosine Kinase | 7,5 |
| KN-62 | Ca2+/calmodulin-dependent pro | Others | 7,5 |
| KN-93 Phosphate | CaMK | Others | 7,5 |
| KRN 633 | VEGFR, PDGFR | Protein Tyrosine Kinase | 7,5 |
| KU-0063794 | mTOR | PI3K/Akt/mTOR | 7,5 |
| KU-55933 (ATM Kinase Inhibitor | ATM | Others | 7,5 |
| KU-60019 | ATM | DNA Damage | 7,5 |
| KW-2449 | Flt, Bcr-Abl, Aurora Kinase | Angiogenesis | 7,5 |
| KX2-391 | Src | Angiogenesis | 7,5 |
| KY02111 | Wnt/beta-catenin | Stem Cells & Wnt | 7,5 |
| Lapatinib | EGFR, HER2 | Protein Tyrosine Kinase | 7,5 |
| Lapatinib (GW-572016) Ditosyla | EGFR, HER2 | Protein Tyrosine Kinase | 7,5 |
| LDC000067 | CDK | Cell Cycle | 7,5 |
| LDE225 (NVP-LDE225,Erismod | Smoothened | Stem Cells & Wnt | 7,5 |
| LDK378 | ALK | Protein Tyrosine Kinase | 7,5 |
| LDN-214117 | TGF-beta/Smad | TGF-beta/Smad | 7,5 |
| Lenvatinib (E7080) | VEGFR | Protein Tyrosine Kinase | 7,5 |

| | | | |
|---|---|---|---|
| LFM-A13 | BTK | Angiogenesis | 7,5 |
| LGK-974 | Wnt/beta-catenin | Stem Cells & Wnt | 7,5 |
| Linifanib (ABT-869) | PDGFR, VEGFR | Protein Tyrosine Kinase | 7,5 |
| LJH685 | S6 Kinase | PI3K/Akt/mTOR | 7,5 |
| LJI308 | S6 Kinase | PI3K/Akt/mTOR | 7,5 |
| Losmapimod (GW856553X) | p38 MAPK | MAPK | 7,5 |
| LY2090314 | GSK-3 | PI3K/Akt/mTOR | 7,5 |
| LY2157299 | TGF-beta/Smad | TGF-beta/Smad | 7,5 |
| LY2603618 | Chk | Cell Cycle | 7,5 |
| LY2784544 | JAK | JAK/STAT | 7,5 |
| LY2784544 | JAK | JAK/STAT | 7,5 |
| LY2811376 | 5-alpha Reductase | Proteases | 7,5 |
| LY2835219 | CDK | Cell Cycle | 7,5 |
| LY294002 | PI3K | Others | 7,5 |
| LY3023414 | Akt | PI3K/Akt/mTOR | 7,5 |
| LY411575 | Gamma-secretase | Proteases | 7,5 |
| Masitinib (AB1010) | c-Kit, PDGFR, FGFR, FAK | Others | 7,5 |
| MEK162 (ARRY-162, ARRY-43 | MEK | MAPK | 7,5 |
| MGCD-265 | c-Met, VEGFR, Tie-2 | Protein Tyrosine Kinase | 7,5 |
| Milciclib (PHA-848125) | CDK | Cell Cycle | 7,5 |
| MK-0752 | Gamma-secretase | Proteases | 7,5 |
| MK-2206 2HCl | Akt | Others | 7,5 |
| MK-2461 | c-Met, FGFR, PDGFR | Protein Tyrosine Kinase | 7,5 |
| MK-5108 (VX-689) | Aurora Kinase | Cell Cycle | 7,5 |
| MK-8745 | Aurora Kinase | Cell Cycle | 7,5 |
| MLN2480 | Raf | MAPK | 7,5 |
| MLN8054 | Aurora Kinase | Others | 7,5 |
| Motesanib Diphosphate (AMG-7 | VEGFR, PDGFR, c-Kit | Protein Tyrosine Kinase | 7,5 |
| Mubritinib (TAK 165) | HER2 | Protein Tyrosine Kinase | 7,5 |
| Nilotinib (AMN-107) | Bcr-Abl | Angiogenesis | 7,5 |
| Nintedanib (BIBF 1120)_uncerta | VEGFR, PDGFR, FGFR | Protein Tyrosine Kinase | 7,5 |
| NSC 23766 | Rac | Cell Cycle | 7,5 |
| NU6027 | CDK | Cell Cycle | 7,5 |
| NVP-AEW541 | IGF-1R | Protein Tyrosine Kinase | 7,5 |
| NVP-BHG712 | VEGFR, Src, Raf, Bcr-Abl | Protein Tyrosine Kinase | 7,5 |
| NVP-BSK805 2HCl | JAK | JAK/STAT | 7,5 |
| NVP-BVU972 | c-Met | Protein Tyrosine Kinase | 7,5 |
| Oclacitinib | JAK | JAK/STAT | 7,5 |
| Olmutinib (HM61713, BI 148269 | EGFR | Protein Tyrosine Kinase | 7,5 |
| ONO-4059 | BTK | Angiogenesis | 7,5 |
| OSI-027 | mTOR | PI3K/Akt/mTOR | 7,5 |
| OSI-420 | EGFR | Protein Tyrosine Kinase | 7,5 |
| OSI-906 (Linsitinib) | IGF-1R | Others | 7,5 |
| OSI-930 | c-Kit, VEGFR | Protein Tyrosine Kinase | 7,5 |
| OSU-03012 (AR-12) | PDK-1 | Others | 7,5 |
| P276-00 | CDK | Cell Cycle | 7,5 |
| Pacritinib (SB1518) | JAK | JAK/STAT | 7,5 |
| Palomid 529 (P529) | mTOR | PI3K/Akt/mTOR | 7,5 |
| Pazopanib | VEGFR | Protein Tyrosine Kinase | 7,5 |
| Pazopanib HCl | VEGFR, PDGFR, c-Kit | Protein Tyrosine Kinase | 7,5 |
| PD0325901 | MEK | DNA Damage | 7,5 |
| PD168393 | EGFR | Protein Tyrosine Kinase | 7,5 |
| PD173074 | FGFR, VEGFR | Angiogenesis | 7,5 |
| PD173955 | Bcr-Abl | Angiogenesis | 7,5 |
| PD184352 (CI-1040) | MEK | MAPK | 7,5 |
| PD318088 | MEK | MAPK | 7,5 |
| PD98059 | MEK | MAPK | 7,5 |
| Pelitinib (EKB-569) | EGFR | Protein Tyrosine Kinase | 7,5 |
| PF-00562271 | FAK | Angiogenesis | 7,5 |
| PF-04217903 | c-Met | Others | 7,5 |
| PF-04691502 | mTOR, PI3K, Akt | PI3K/Akt/mTOR | 7,5 |
| PF-3758309 | PAK | Cytoskeletal Signaling | 7,5 |
| PF-431396 | FAK | Angiogenesis | 7,5 |
| PF-4708671 | S6 Kinase | PI3K/Akt/mTOR | 7,5 |
| PF-477736 | Chk | Cell Cycle | 7,5 |
| PF-5274857 | Hedgehog/Smoothened | Stem Cells & Wnt | 7,5 |
| PF-543 | SphK1 | GPCR & G Protein | 7,5 |
| PF-562271 | FAK | Angiogenesis | 7,5 |
| PF-573228 | FAK | Angiogenesis | 7,5 |
| PFK15 | 6-phosphofructo-2-kinase (PFK | Others | 7,5 |
| PH-797804 | p38 MAPK | MAPK | 7,5 |
| PHA-665752 | c-Met | Others | 7,5 |
| PHA-680632 | Aurora Kinase | Cell Cycle | 7,5 |
| PHA-767491 | CDK | Cell Cycle | 7,5 |
| PHA-793887 | CDK | Cell Cycle | 7,5 |
| Phenformin HCl | AMPK | PI3K/Akt/mTOR | 7,5 |
| PHT-427 | Akt, PDK-1 | PI3K/Akt/mTOR | 7,5 |
| PI-103 | DNA-PK, PI3K, mTOR | Neuronal Signaling | 7,5 |

| | | | |
|---|---|---|---|
| Piceatannol | Syk | Angiogenesis | 7,5 |
| PIK-293 | PI3K | PI3K/Akt/mTOR | 7,5 |
| PIK-294 | PI3K | PI3K/Akt/mTOR | 7,5 |
| PIK-93 | PI3K, VEGFR | PI3K/Akt/mTOR | 7,5 |
| Pimasertib (AS-703026) | MEK | MAPK | 7,5 |
| Pirfenidone | TGF-beta/Smad | TGF-beta/Smad | 7,5 |
| PLX-4720 | Raf | Others | 7,5 |
| PLX7904 | Raf | MAPK | 7,5 |
| Ponatinib (AP24534) | Bcr-Abl, VEGFR, FGFR, PDGF | Angiogenesis | 7,5 |
| PP1 | Src | Angiogenesis | 7,5 |
| PP121 | DNA-PK, mTOR, PDGF | Protein Tyrosine Kinase | 7,5 |
| PP2 | Src | Angiogenesis | 7,5 |
| PP242 | mTOR | PI3K/Akt/mTOR | 7,5 |
| PQ 401 | IGF-1R | Protein Tyrosine Kinase | 7,5 |
| PRI-724 | Wnt/beta-catenin | Stem Cells & Wnt | 7,5 |
| PRT062607 (P505-15, BIIB057) | Syk | Angiogenesis | 7,5 |
| Purvalanol A | CDK | Cell Cycle | 7,5 |
| Quercetin | PI3K, PKC, Src, Sirtuin | Epigenetics | 7,5 |
| Quizartinib (AC220) | Flt | Angiogenesis | 7,5 |
| R406 | Syk, Flt | Angiogenesis | 7,5 |
| R406 (free base) | Syk | Angiogenesis | 7,5 |
| R547 | CDK | Cell Cycle | 7,5 |
| RAF265 (CHIR-265) | Raf, VEGFR | MAPK | 7,5 |
| Rapamycin (Sirolimus) | mTOR | DNA Damage | 7,5 |
| Refametinib (RDEA119, Bay 86- | MEK | Others | 7,5 |
| Regorafenib (BAY 73-4506) | c-Kit, Raf, VEGFR | Protein Tyrosine Kinase | 7,5 |
| RepSox | TGF-beta/Smad | TGF-beta/Smad | 7,5 |
| Ridaforolimus (Deforolimus, MK- | mTOR | PI3K/Akt/mTOR | 7,5 |
| Rigosertib (ON-01910) | PLK | Cell Cycle | 7,5 |
| RKI-1447 | ROCK | Cell Cycle | 7,5 |
| RN486 | BTK | Angiogenesis | 7,5 |
| Ro 31-8220 Mesylate | PKC | TGF-beta/Smad | 7,5 |
| Ro-3306 | CDK | Cell Cycle | 7,5 |
| Ro3280 | PLK | Cell Cycle | 7,5 |
| RO4929097 | Y-Secretase | Proteases | 7,5 |
| Roscovitine (Seliciclib,CYC202) | CDK | Others | 7,5 |
| Ruxolitinib (INCB018424) | JAK | JAK/STAT | 7,5 |
| S-Ruxolitinib (INCB018424) | JAK | JAK/STAT | 7,5 |
| SANT-1 | Smoothened | Stem Cells & Wnt | 7,5 |
| SAR131675 | VEGFR | Protein Tyrosine Kinase | 7,5 |
| SAR245409 (XL765) | PI3K, mTOR | PI3K/Akt/mTOR | 7,5 |
| Saracatinib (AZD0530) | Src, Bcr-Abl | Angiogenesis | 7,5 |
| SB202190 (FHPI) | p38 MAPK | PI3K/Akt/mTOR | 7,5 |
| SB203580 | p38 MAPK | Transmembrane Transpo | 7,5 |
| SB216763 | GSK-3 | Others | 7,5 |
| SB239063 | p38 MAPK | MAPK | 7,5 |
| SB415286 | GSK-3 | PI3K/Akt/mTOR | 7,5 |
| SB431542 | TGF-beta/Smad | Others | 7,5 |
| SB505124 | TGF-beta/Smad | TGF-beta/Smad | 7,5 |
| SB525334 | TGF-beta/Smad | TGF-beta/Smad | 7,5 |
| SB590885 | Raf | MAPK | 7,5 |
| SC-514 | I_B/IKK | NF-_B | 7,5 |
| SC1 | ERK | MAPK | 7,5 |
| Schisandrin B (Sch B) | ATM/ATR | PI3K/Akt/mTOR | 7,5 |
| Selumetinib (AZD6244) | MEK | MAPK | 7,5 |
| Semagacestat (LY450139) | Gamma-secretase | Proteases | 7,5 |
| Semaxanib (SU5416) | VEGFR | Protein Tyrosine Kinase | 7,5 |
| SGI-1776 free base | Pim | JAK/STAT | 7,5 |
| SGI-7079 | VEGFR | Protein Tyrosine Kinase | 7,5 |
| SH-4-54 | STAT | JAK/STAT | 7,5 |
| Skepinone-L | p38 MAPK | MAPK | 7,5 |
| SKI II | sphingosine kinase (SphK) | GPCR & G Protein | 7,5 |
| SL-327 | MEK | Others | 7,5 |
| SMI-4a | Pim | JAK/STAT | 7,5 |
| SNS-032 (BMS-387032) | CDK | Others | 7,5 |
| SNS-314 Mesylate | Aurora Kinase | Others | 7,5 |
| Sorafenib | Raf | MAPK | 7,5 |
| Sotrastaurin | PKC | TGF-beta/Smad | 7,5 |
| SP600125 | JNK | MAPK | 7,5 |
| SSR128129E | FGFR | Angiogenesis | 7,5 |
| STA-21 | STAT | JAK/STAT | 7,5 |
| SU11274 | c-Met | Neuronal Signaling | 7,5 |
| SU6656 | Src | Angiogenesis | 7,5 |
| SU9516 | CDK | Cell Cycle | 7,5 |
| Sunitinib Malate | VEGFR, PDGFR, c-Kit, Flt | Microbiology | 7,5 |
| TAE226 (NVP-TAE226) | FAK | Angiogenesis | 7,5 |
| TAK-285 | EGFR, HER2 | Protein Tyrosine Kinase | 7,5 |
| TAK-632 | Raf | MAPK | 7,5 |

| | | | |
|---|---|---|---|
| TAK-715 | p38 MAPK | MAPK | 7,5 |
| TAK-733 | MEK | MAPK | 7,5 |
| TAK-901 | Aurora Kinase | Cell Cycle | 7,5 |
| Taladegib (LY2940680) | Hedgehog,Hedgehog/Smoothen | Stem Cells & Wnt | 7,5 |
| TCS 359 | FLT3 | Angiogenesis | 7,5 |
| TDZD-8 | GSK-3 | PI3K/Akt/mTOR | 7,5 |
| Telatinib | VEGFR, PDGFR, c-Kit | Protein Tyrosine Kinase | 7,5 |
| Temsirolimus (CCI-779, NSC 6 | mTOR | Neuronal Signaling | 7,5 |
| Tepotinib (EMD 1214063) | c-Met | Protein Tyrosine Kinase | 7,5 |
| TG003 | CDK | Cell Cycle | 7,5 |
| TG100-115 | PI3K | PI3K/Akt/mTOR | 7,5 |
| TG101209 | Flt, JAK, c-RET | JAK/STAT | 7,5 |
| TG101348 (SAR302503) | JAK | JAK/STAT | 7,5 |
| TGX-221 | PI3K | PI3K/Akt/mTOR | 7,5 |
| Theophylline | TGF-beta/Smad | TGF-beta/Smad | 7,5 |
| Thiazovivin | ROCK | Cell Cycle | 7,5 |
| TIC10 Analogue | Akt | PI3K/Akt/mTOR | 7,5 |
| Tideglusib | GSK-3 | PI3K/Akt/mTOR | 7,5 |
| Tie2 kinase inhibitor | Tie-2 | Protein Tyrosine Kinase | 7,5 |
| Tivantinib (ARQ 197) | c-Met | Protein Tyrosine Kinase | 7,5 |
| Tivozanib (AV-951) | VEGFR, c-Kit, PDGFR | Protein Tyrosine Kinase | 7,5 |
| Tofacitinib (CP-690550,Tasociti | JAK | JAK/STAT | 7,5 |
| Tofacitinib (CP-690550) Citrate | JAK | JAK/STAT | 7,5 |
| Torin 2 | mTOR | PI3K/Akt/mTOR | 7,5 |
| TPCA-1 | IKK | NF-_B | 7,5 |
| Trametinib (GSK1120212) | MEK | MAPK | 7,5 |
| Triciribine | Akt | Others | 7,5 |
| TSU-68 (SU6668, Orantinib) | VEGFR, PDGFR , FGFR | Protein Tyrosine Kinase | 7,5 |
| TWS119 | GSK-3 | PI3K/Akt/mTOR | 7,5 |
| TWS119 | GSK-3 | PI3K/Akt/mTOR | 7,5 |
| Tyrphostin 9 | EGFR | Protein Tyrosine Kinase | 7,5 |
| Tyrphostin AG 1296 | PDGFR | Protein Tyrosine Kinase | 7,5 |
| Tyrphostin AG 879 | HER2 | Protein Tyrosine Kinase | 7,5 |
| U0126-EtOH | MEK | Others | 7,5 |
| Ulixertinib (BVD-523, VRT75227 | ERK | MAPK | 7,5 |
| Uprosertib (GSK2141795) | Akt | PI3K/Akt/mTOR | 7,5 |
| URMC-099 | Abl1; LRRK2; MLK3; MLK1 | Others | 7,5 |
| Vacquinol-1 | JNK | MAPK | 7,5 |
| Varlitinib | EGFR | Protein Tyrosine Kinase | 7,5 |
| Vatalanib (PTK787) 2HCl | VEGFR, c-Kit, Flt | Others | 7,5 |
| VE-821 | ATM/ATR | DNA Damage | 7,5 |
| VE-822 | ATM/ATR | PI3K/Akt/mTOR | 7,5 |
| Vemurafenib (PLX4032, RG720 | Raf | MAPK | 7,5 |
| Vismodegib (GDC-0449) | Hedgehog, P-gp | Neuronal Signaling | 7,5 |
| Volasertib (BI 6727) | PLK | Cell Cycle | 7,5 |
| VPS34-IN1 | PI3K | PI3K/Akt/mTOR | 7,5 |
| VS-5584 (SB2343) | PI3K | PI3K/Akt/mTOR | 7,5 |
| VX-11e | ERK | MAPK | 7,5 |
| VX-680 (Tozasertib, MK-0457) | Aurora Kinase | Endocrinology & Hormone | 7,5 |
| VX-702 | p38 MAPK | MAPK | 7,5 |
| VX-745 | p38 MAPK | MAPK | 7,5 |
| WAY-600 | mTOR | PI3K/Akt/mTOR | 7,5 |
| WH-4-023 | Src | Angiogenesis | 7,5 |
| WHI-P154 | JAK, EGFR | JAK/STAT | 7,5 |
| WIKI4 | Wnt/beta-catenin | Stem Cells & Wnt | 7,5 |
| Wnt agonist 1 | Wnt/beta-catenin | Stem Cells & Wnt | 7,5 |
| Wnt-C59 (C59) | Wnt/beta-catenin | Stem Cells & Wnt | 7,5 |
| WP1066 | JAK | JAK/STAT | 7,5 |
| WYE-125132 (WYE-132) | mTOR | PI3K/Akt/mTOR | 7,5 |
| WYE-354 | mTOR | PI3K/Akt/mTOR | 7,5 |
| WZ3146 | EGFR | Protein Tyrosine Kinase | 7,5 |
| WZ4002 | EGFR | Protein Tyrosine Kinase | 7,5 |
| WZ4003 | AMPK | PI3K/Akt/mTOR | 7,5 |
| WZ8040 | EGFR | Protein Tyrosine Kinase | 7,5 |
| XAV-939 | Wnt/beta-catenin | Stem Cells & Wnt | 7,5 |
| XL019 | JAK | JAK/STAT | 7,5 |
| XMD8-92 | ERK | MAPK | 7,5 |
| Y-27632 2HCl | ROCK | Others | 7,5 |
| YM201636 | PI3K | PI3K/Akt/mTOR | 7,5 |
| YO-01027 | Gamma-secretase | Proteases | 7,5 |
| ZM 306416 | VEGFR | Protein Tyrosine Kinase | 7,5 |
| ZM 323881 HCl | VEGFR | Protein Tyrosine Kinase | 7,5 |
| ZM 336372 | Raf | MAPK | 7,5 |
| ZM 39923 HCl | JAK | JAK/STAT | 7,5 |
| ZM 447439 | Aurora Kinase | Others | 7,5 |
| Zotarolimus(ABT-578) | mTOR | PI3K/Akt/mTOR | 7,5 |
| ZSTK474 | PI3K | Neuronal Signaling | 7,5 |
